# The effect of dispersal and preferential mating on the genetic control of mosquitoes

**DOI:** 10.1101/2020.05.25.114413

**Authors:** Doran Khamis, Claire El Mouden, Klodeta Kura, Michael B. Bonsall

## Abstract

Mosquito-borne diseases cause significant social and economic damage across much of the globe. New biotechnologies that utilise manipulations of the mosquito genome have been developed to combat disease. The successful implementation of genetic mosquito control technologies may depend upon ecological, evolutionary and environmental factors, as well as the specifications of the chosen technology. Understanding the influence of these external factors will help inform how best to deploy a chosen technology to control vectors of infectious diseases. We use a continuous-time stochastic spatial network model of a mosquito life-cycle coupled to population genetics models to investigate the impact of releasing seven types of genetic control technology: a self-limiting lethal gene, two underdominance threshold gene drives, two homing gene drives and two *Wolbachia* systems. We apply the mathematical framework to understand control interventions of two archetypes of mosquito species: a short-range dispersing *Aedes aegypti* and comparatively longer-range dispersing *Anopheles gambiae*. We show that mosquito dispersal behaviour is an extremely important factor in determining the outcome of a release programme. Assortative mating – where the mating success of genetically modified males is lower than their wild counterparts – can facilitate the spatial containment of gene drives. The rapid evolution of strong mating preference can damage the efficacy of control efforts for all control technologies. We suggest that there cannot be a one-size-fits-all approach to regulation and implementation of vector control; there must be application-specific control plans that take account of understudied ecological, evolutionary and environmental factors.

## 1 Introduction

Mosquito-borne diseases pose a major threat to the health and economies of societies around the world. More than 80% of the world’s population live in areas at risk from a major vector-borne disease such as malaria, dengue or Zika [69]. Due to increasing insecticide and drug resistance, urbanization and climate change, the burden of many vector-borne diseases has increased in recent years. In 2016 alone, Zika cost the Americas’ poorest tourism-based economies a devastating $3.5B [73]. In response to this social and economic crisis, a range of new biotechnologies are being developed that seek to disrupt vector populations. These either aim to suppress the mosquito population or to change the genetic makeup of the wild population so as to render it unable to transmit disease [2]. These technologies fall into two categories: self-limiting – so called because their effects disappear when releases stop, as the genetic components are lost from the population – and self-sustaining – intended to persist indefinitely in the target population after release, possibly increasing in frequency and spreading – which can be further subdivided into threshold drives, homing drives and naturally occuring proliferative elements like the intracellular bacteria *Wolbachia* (that spread through the mechanism of cytoplasmic incompatibility). Self-limiting technologies are undergoing field trials [10, 28], while self-sustaining genetic technologies are still years from field testing [although *Wolbachia* is already being used with success in the field, 60]. A fundamental problem for regulators and public health agencies is that the novelty of these technologies means the potential benefits and risks are only beginning to be understood. A key concern is how these technologies will spread between mosquito populations in space and time. More generally, a better understanding of the spatial response of specific vector populations to population suppression attempts is essential to identify optimal release strategies and important parameters governing release outcomes. In the absence of trial data, empirically-based models that account for species-specific ecology must help guide research, regulation and implementation.

Research by behavioural biologists and entomologists has shown that the spatial ecology of mosquitoes is strongly species-specific. This is pertinent for predictions of the performance of control efforts in terms of spread or containment. *Aedes aegypti* mosquitoes are known to travel only short distances over their lifetime, with a large percentage living within a single house or moving between neighbouring houses very infrequently [32, 34]. Conversely, *Anopheles gambiae* mosquitoes regularly move between neighbouring villages [64, 66], and recent research has shown that they traverse large distances to repopulate arid regions at the end of the dry season [15]. The difference in these dispersal behaviours must be taken into account when deciding how best to design and implement genetic control technologies.

Many studies have investigated the effect of genetic mosquito control technologies within a single species from the perspective of spatial spread or containment. The most complex and richly parameterised of these used agent-based stochastic modelling approaches, capturing explicit spatial heterogeneity through the distribution of larval breeding sites [48, 59], blood-feeding sites [56] and bodies of standing water [58]. The most simple and intuitive approaches consider the growth and decline in frequency of invading alleles in population genetics frameworks [17, 50]. These complementary modelling approaches have allowed predictions of the most effective spatial release strategies for various control technologies [38, 46] when controlling *Aedes aegypti* populations. Other studies have investigated the necessary parameters for success of gene drive technologies in spatially structured *Anopheles gambiae* populations [22, 56]. Marshall [49] undertook a thorough risk assessment of current control technologies through the lens of the probability of escape, survival and spread from an ambient field cage, and abstracted to multiple species of mosquito—however, the theoretical treatment used did not allow modelling of explicit spatial structure, with a later paper suggesting the need for characterization of local ecology before control strategies are implemented [50]. In the current study, we choose to take aim with our theoretical approach at the ground between complex agent-based models and simple population genetic models by utilising a network approach [extended from that developed in 75] to investigate gene flow between and the effect of control technologies on a mosquito metapopulation. Crucially, our goal is to bridge the gap between mosquito species by explicitly comparing control technologies for two mosquito archetypes: a short-range dispersing species (like *Aedes aegypti*) and a comparatively longer-range dispersing species (like *Anopheles gambiae*).

In this work we compare spatial models of seven control technologies via the metric of their efficacy for two different mosquito species archetypes. The technologies are: a late-acting lethal gene self-limiting control [1], where transgenic males are released whose offspring die at a late stage of larval development; two types of engineered underdominance threshold gene drive [16] (a homologous single-locus technology and a non-homologous system using two loci), which exploit bi-stable gene frequencies (due to heterozygote fitness disadvantage) to drive introduced genes to fixation once their gene frequencies pass a threshold in the population; two types of homing gene drive [such as those proposed using CRISPR/Cas9 nucleases 26], that uses dynamic gene editing in the germline to drive an introduced gene through a population at a greater than Mendelian rate (one version in which the drive gene and lethal payload are combined and one version where the drive component is inherited separately from the lethal gene); and two types of *Wolbachia*, unidirectional (one bacteria strain) and bidirectional (two strains), which are a natural maternally inherited drive system causing offspring survival bias through cytoplasmic incompatibility. There exists no clear comparison of these categories of vector control across different mosquito species within a consistent ecological and population genetic framework. Further to this, there are several key ecological, evolutionary and environmental factors that are still poorly understood and are only recently starting to receive deserved attention. Our study focuses on how dispersal and mating behaviour interact with imposed fitness costs to impact control efforts using the seven technologies listed above.

There is scientific consensus that populations may evolve resistance to these genetic technologies [7, 55]. However, it is unknown if resistance could result from selection for greater female choosiness and if it can, whether it could evolve rapidly enough to impair the effectiveness of a control effort. At release, mass-reared males will likely exhibit quite obvious phenotypic differences from the wild strain, so if this ‘behavioural resistance’ evolves rapidly, females may be able to discriminate between mass reared and wild type. Conversely, if female preference has only a weak influence on mate choice, then male competition will be the phenomenon driving mating behaviour, and if released males are less competitive than wild males a different sort of behavioural resistance will manifest. Behavioural resistance has been observed in many dipteran control programs including screwworm [9], melon fly [35], and medfly [52]; it can appear within a few generations, and has led to control programs being abandoned [52]. The probability of behavioural resistance developing and the potential challenge that it could present for the successful application of genetic control in mosquitoes is unknown. Both possibilities – female choosiness and male competitiveness – will impact on the mating success of genetically modified males with wild females, and we investigate this here.

Here, we use coupled continuous-time ecological and population genetics models on a spatial network to compare the impact of a self-limiting technology, underdominance threshold drives, homing-based gene drives and *Wolbachia* drives on wild mosquito populations of two distinct archetypal species. We investigate conditions for successful suppression or replacement, either global or contained, across a network of randomly generated villages (or neighbourhoods) under the influence of changing ecological, evolutionary and genetic pressures.

## 2 Methods

We use a stage-structured population model for the mosquito, taking into account an aquatic juvenile stage (*B*) and an adult stage (*N*). The dynamics are captured by the following set of ordinary differential equations:

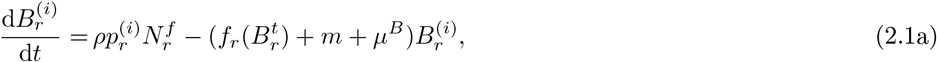

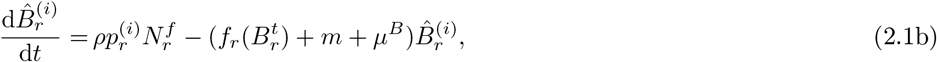

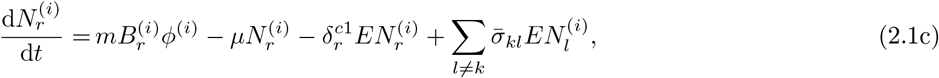

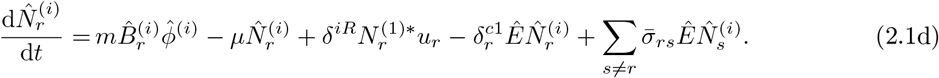

A superscript “(*i*)” denotes genotype; a subscript “*r*” denotes the spatial site (or network node); a hat denotes males or male-specific quantities. Thus, 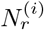 represents adult females of genotype *i* at node *r* and 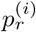 is the proportion of offspring of genotype *i* at node *r* that arise from the genetic mating crosses (we assume an equal sex ratio for all offspring). In (2.1), *ρ* is the oviposition rate of the adult females, *m* is the rate at which larvea mature into adults and *μ* and *μ*^*B*^ are the density-independent mortality rates for adults and juveniles, respectively. For simplicity, we assume that these life history parameters (*ρ*, *m*, *μ* and *μ*^*B*^) are homogeneous across spatial sites and genotype (though this will not be the case in general, and relaxing this assumption would allow an interesting investigation of heterogeneity to be performed). The Kronecker delta *δ*^*iR*^ ensures released mosquitoes are of the correct genotype, labelled “*R*” here. The number of mosquitoes released per day is *u*_*r*_. The sum of all male and female larvae is

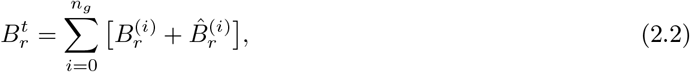

where *n*_*g*_ is the number of genotypes (which is technology dependent), while the sum of adult females is 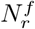. The Kronecker delta 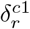 is equal to one (*c* = 1) if connections to other nodes from node *r* are open and zero (*c* ≠ 1) if node *r* is unconnected (movement of mosquitoes is discussed further below and in appendix S1a). Parameter values and definitions are listed in table 1.

**Table 1.** Ecological model and release strategy parameter definitions and values for the two archetypal mosquito species.

| Symbol | Description | *Aedes* | *Anopheles* | Notes |
| --- | --- | --- | --- | --- |
| $\rho$ | daily per adult female oviposition rate | 16 | $\frac{52}{3}$ | [20, 62, 63] |
| $m$ | daily larval maturation rate | $-\ln \frac{13}{15}$ | $-\ln \frac{9}{10}$ | [6, 36, 45] |
| $\mu$ | daily adult mosquito death rate | $-\ln \frac{88}{100}$ | $-\ln 0.925$ | [13, 18, 21, 24, 25, 44, 53, 54] |
| $\mu_B$ | daily density-independent larval death rate | $-\ln 0.985$ | $-\ln 0.9805$ | [39, 43, 45] |
| $\eta$ | strength of density dependence | 0.9 | 0.9 | $\eta < 1$ implies contest-type density dependence |
| $\nu$ | scale of larval density-dependence | eq. (2.4), (S1.6) | eq. (2.4), (S1.6) | regulates seasonal population fluctuations |
| $k^*$ | Average vector-to-host ratio | 15 | - | Undergoes seasonal and stochastic variation, see appendix S1b |
| $H$ | Nominal host population per node | 4 | - | We do not model host-vector interaction, this just sets the scale of the computations |
| $u$ | Weekly release size | 2000 | - | Spread over four nodes of Village 1, see appendix S2 |
| $c_1$ | Ambient fitness cost of carrying a single modified allele | 0.05 | - | Fitness costs combine multiplicatively (appendix S2) |
| $c_2$ | Female-specific lethality efficiency of carrying single lethal allele | 0.85 | - | Fitness costs combine multiplicatively (appendix S2) |

The fitness costs *φ*^(*i*)^ and 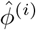, which may differ, act at the end of the pupal stage, allowing all larvae to compete for resources in the larval habitat. This reduces the risk of unintended population rebound [over-compensatory density dependence acting to increase the reproductive fitness of a cohort maturing in a less competitive environment 4, 74]. Fitness costs imposed by modified alleles (either intended lethal payloads or the ambient burden of carrying a transgene) combine multiplicatively (aside from when combinations of transgenes suppress lethal effects, as in the underdominance systems). For instance, work has shown that density-dependent effects can have important implications for the spread of *Wolbachia* in mosquitoes [30, 31]. The density dependence experienced by larvae at node *r* is given by [51]

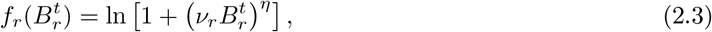

where *ν*_*r*_ is inversely proportional to the carrying capacity at node *r* and *η* governs the approach rate to the carrying capacity. The wild-type carrying capacity is defined through the vector-to-host ratio 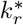 and a nominal host population *H* per node: 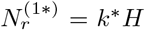 (with the superscript * denoting the equilibirum value). The human population *H* is assumed constant and equal at all nodes, though in this model *H* acts purely as a multiplicative constant in the density dependent regulation (i.e. we are not implying a mechanistic relationship that causes *H* to limit the vector population; *H* is just used as a mathematical convenience to set the scale for the simulations). The density dependence parameters act to enforce this equilibrium population through the definition

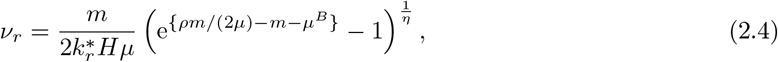

which holds separately at each node *r* (possibly producing different values of *ν*_*r*_). Seasonality and localised environmental stochasticity (node-to-node heterogeneity) is captured by changing 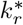 stochastically and periodically (using a noisy sinusoidal signal bounded above a small value, see appendix S1b).

Movement of adult mosquitoes between nodes (described by the movement weighting matrix 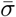 in (2.1)) is treated stochastically, with connections between pairs of nodes opening and closing each day as governed by a Gaussian probability distribution dependent on the distance between the nodes [see appendix S1a and 75]. The opening and closing of connections between nodes can be thought of as capturing physical events such as the closing of a window or door between neighbouring nodes or a strong wind or heavy shower preventing travel between distant nodes. The distribution of migrants from a given node among all the available connections is governed by the species-specific Laplacian dispersal kernels (appendix S1a). A percentage of the vector population at a given home site will move each day when at least one connection to a target site is open, with that percentage sampled randomly for each node on each day from a fixed Gaussian distribution. The variation in distance from a home site to multiple valid target sites governs how the migrants are spread between the target sites, with closer sites receiving more migrants. The distance between sites also dictates whether the path between the sites will be open. If no paths are open, no migration occurs and the dispersal rate is (temporarily) zero. Shorter paths are more likely to be open, meaning dispersal to close targets occurs at a higher rate. The number of open connections per node is capped at thirty.

The action of each genetic control technology is captured in the population genetics model, which accounts for homing rates (the efficiency with which a gene drive heterozygote is converted into a gene drive homozygote), lethality efficiencies (of the genetic construct designed to cause population suppression), unintended fitness costs (of carrying a genetic modification) and female wild-type mating preferences. The number of possible genotypes is dictated by the control technology chosen (see appendix S3). Mating preference of wild-type females is defined as the fraction of females that, upon coming into contact with a male of a non-wild genotype (homozygotes or heterozygotes of modified alleles), will instead mate with a wild (homozygous) male (see appendix S3a). The mechanism for the preference is not assumed: the resulting skew in offspring proportion could be achieved, for example, through explicit female choice (phenotypic preference); through a reduced ability of genetically modified males to compete for mates with wild males; or due to a combination of these (or other) factors. We assume each control technology results in female-specific lethality (allowing for the unique effect of *Wolbachia*), but also imposes an unintended, ambient fitness cost which is borne by those mosquitoes of either sex that carry the transgene.

Simulations span three years, with control implemented after the first year to allow the system to reach its natural equilibrium. The spatial structure of the network (shown in Supplementary Information fig. S1) consists of a square domain of area 5km^2^ containing three villages (or neighbourhoods) of fifty houses (nodes, acting as mosquito breeding sites) each (Village 2 close to Village 1; Village 3 far from Village 1 but close to Village 2), with the positions of the houses randomly generated each time the model is run. The model was written in C++ and source code is available at https://osf.io/4f9jk/.

## 3 Results

We investigate how dispersal, mating success and genetic fitness costs affect the performance of seven population suppression technologies: a self-limiting technology (SL), two underdominance threshold drives – homologous (UD) and non-homologous (NHUD) – two homing gene drives – combined homing and lethal gene (CGD) and separated homing and lethal genes (SGD) – and two cytoplasmic incompatibility drives – unidirectional *Wolbachia* (WBU) and bidirectional *Wolbachia* (WBB). Importantly, we investigate how these results differ between a short-range dispersing species like *Aedes aegypti* (Ae) and a comparatively longer-range dispersing species like *Anopheles gambiae* (An). Also investigated is how the disparate population sizes of the *Anopheles* compared with the *Aedes* mosquito (*Anopheles* can have effective population sizes larger by a factor of many thousands) affects control efforts. Resistance and invasiveness is then studied by simulating the development of behavioural and genetic resistance to control efforts; the invasiveness of *Wolbachia* drives is studied in the context of the possibility of a positive fitness bias.

Fitness costs are modelled as a reduction in the probability that a mosquito will reach reproductive age. We consider two types of fitness cost: intended action and unintended burden. The intended fitness costs are those that result from the desired phenotypic consequences of the control technology (the lethality efficiency) and we assume these only affect females (female-specific lethality) barring the specific action of cytoplasmic incompatibility; the unintended, or ambient, costs arise as a consequence of carrying a modified gene (such as increased mutagenesis or unforeseen phenotypic effects) and are borne equally by both sexes. Often only the lethality efficiency of a technology is studied; we also investigate the effect of the ambient fitness cost for several reasons: (i) it should be possible to adapt the design of genetic technologies to control, at least partly, this parameter; (ii) the relative fitness of the transgenic mosquito is a key parameter governing its establishment and persistence [e.g. 8]; (iii) testing a range of unintended fitness costs hints at the macroscopic effects of the likely build-up of off-target cutting and resulting deleterious mutations when CRISPR/Cas drive systems are used [12, 76]; and (iv) the ambient cost has been shown to be often more important than the lethality efficiency in governing the success of a control strategy [40].

In all following results, we use a release strategy of weekly pulses of 2000 modified males, spread over four equidistant release sites equally spaced between the centre of Village 1 and the village edge (in a square pattern), for one year. (The experiments and reasoning that led to this choice can be found in appendix S2). Releases are then stopped, and the simulations are continued for a year to examine whether and how the vector population rebounds after control is ceased. In the case of the *Wolbachia* systems, both modified males and females are relased by necessity, and releases are spread equally between both sexes in the unidirectional system and between both sexes and both *Wolbachia* strains in the bidirectional system. We study genetic controls designed both for population suppression and replacement. Suppression is the reduction of the abundance of a target population; replacement systems aim to make the vector more refractory to the pathogen they carry. In this study, we use the word “suppression” in the case of the suppressive technologies (self-limiting, homing gene drives) to mean the reduction in abundance of total female numbers irrespective of genotype, while in the case of replacement-focussed technologies (underdominance drives and *Wolbachia*) we mean the reduction in abundance of wild-type females only (the modified females are assumed by the mechanism specific to the system to be ineffective vectors of disease). Results are colour coded green (triangles), blue (squares) and red (squares) to indicate suppression in Village 1, Villages 1 and 2, and Villages 1, 2 and 3, respectively. Further colour coding indicates whether suppression is temporary (light, year of control only) or lasting (dark, both year of control and year after). Gold colouring indicates that suppression was delayed until the year after the control period (for example a gold square denotes suppression in all three villages that takes effect the year after releases cease, while a gold triangle denotes suppression in Village 1 the year after releases cease). Black circles denote simulations wherein no suppression is achieved in any village.

### 3.1 Dispersal

Understanding the dispersal behaviour of a mosquito species is vitally important to be able to predict the spatial spread of a released genetic modification. We simulate control efforts for an “average” mosquito species (all life history parameters are taken as the mean of those of the *Ae. aegypti* and *An. gambiae* listed in table 1), and vary the ‘dispersal level’ of this species, which changes the migration distance probability distribution (appendix S1a). At a dispersal level of 0.25, the migration distance matches that of the *Aedes* archetype; at a dispersal level of 0.75 the migration distance matches that of the *Anopheles* archetype. The dispersal level also changes the probability distribution governing the fraction of the population of a node that migrate each day, and linearly scales between 0% and 15% for dispersal levels in [0, 1].

We find that varying the average migration distance and the percentage of a node’s population that migrates strongly affects whether or not the genetic control can be spatially contained (fig. 1). for dispersal levels lower than around 0.5 (7.5% moving each day, up to around 750m per lifetime), all technologies can be expected to cause local suppression (or replacement) with eventual repopulation from neighbouring villages. For invasive technologies like the CGD, WBU and WBB we see that for slowly dispersing species, global suppression is possible (though perhaps delayed) for low ambient fitness costs as the level of wild-type repopulation is too weak (fig. 1). This transition from global to local suppression at lower dispersal levels is interesting as it occurs in the range of dispersal behaviours shown by *Aedes aegypti* [32, 34], suggesting containment for this species may depend sensitively on the combination of fitness costs imposed by the control technology. For highly dispersive species, it is possible that even underdominance drives may cause global suppression (fig. 2). It is worth highlighting how the SGD, in which the homing gene is inherited at the normal Mendelian rate, is far less invasive than the CGD.

**Figure 1.**
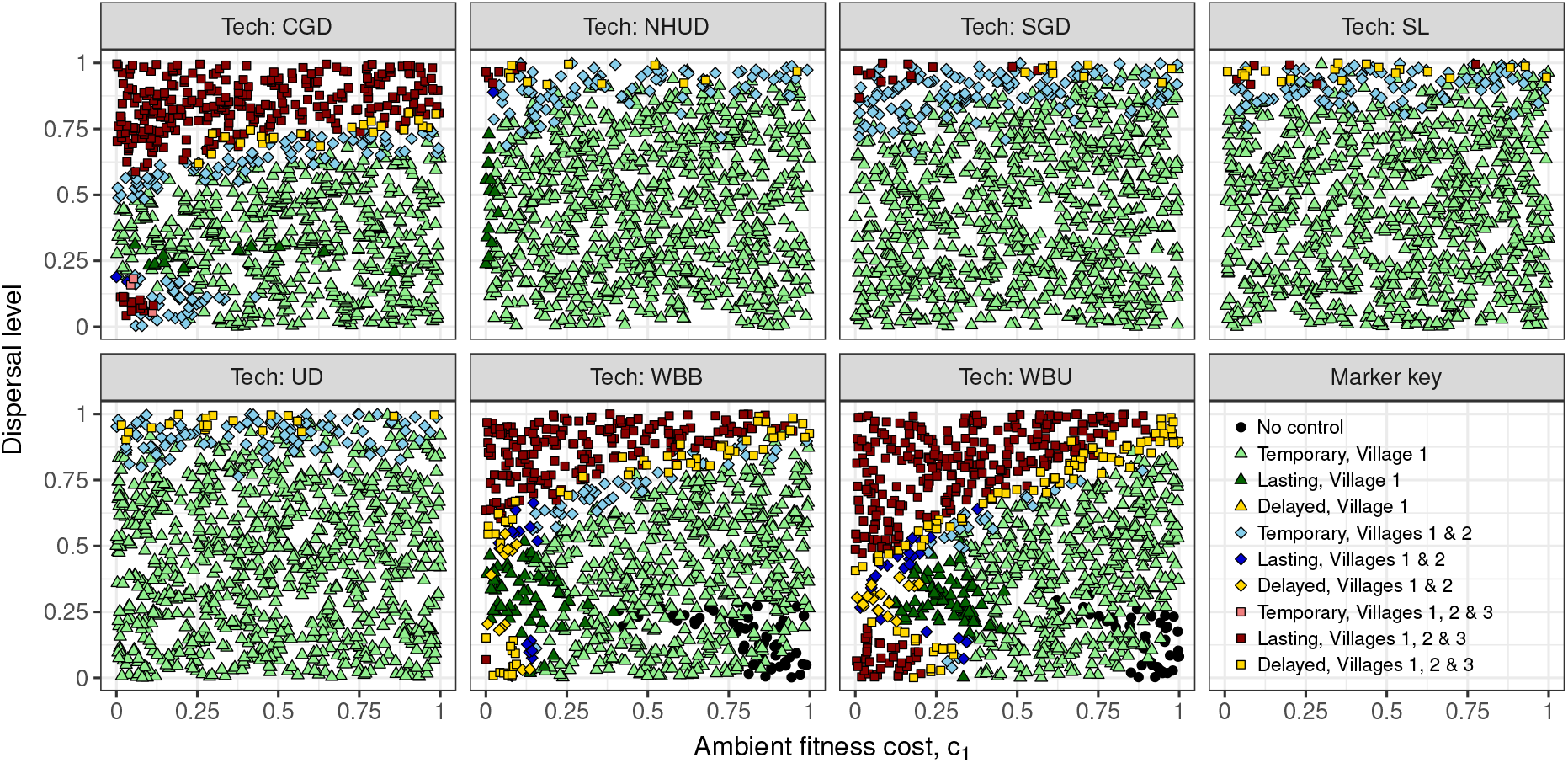
The outcomes of a year-long control effort in one village, with two neighbouring villages, in which 2000 modified mosquitoes were released every week, for varying values of the ambient fitness cost, *c*_1_, imposed by a single modified allele and the dispersal level, which for values in [0, 1] scales the average percentage of mosquitoes migrating from each node each day between 0% and 15%, and scales the probability distribution governing the distances that migrants travel. The lethality efficiency of the payload genes is *c*_2_ = 0.85. The mosquito simulated here is an ‘average’ species, with life history parameters taken as the mean of those listed for *Aedes* and *Anopheles* in table 1. The control technologies are a self-limiting technology (SL), homologous underdominance (UD), non-homologous underdominance (NHUD), a homing gene drive with combined homing and lethal gene (CGD), a homing gene drive with separated homing and lethal genes (SGD), unidirectional single-strain *Wolbachia* (WBU) and bidirectional two-strain *Wolbachia* (WBB). The bottom right panel shows marker shape and colour codings. Temporary control means control in the year of releases but not after releases stop; lasting control means control that continues after releases stop; delayed control means control that does not take affect until the year after releases stop.

**Figure 2.**
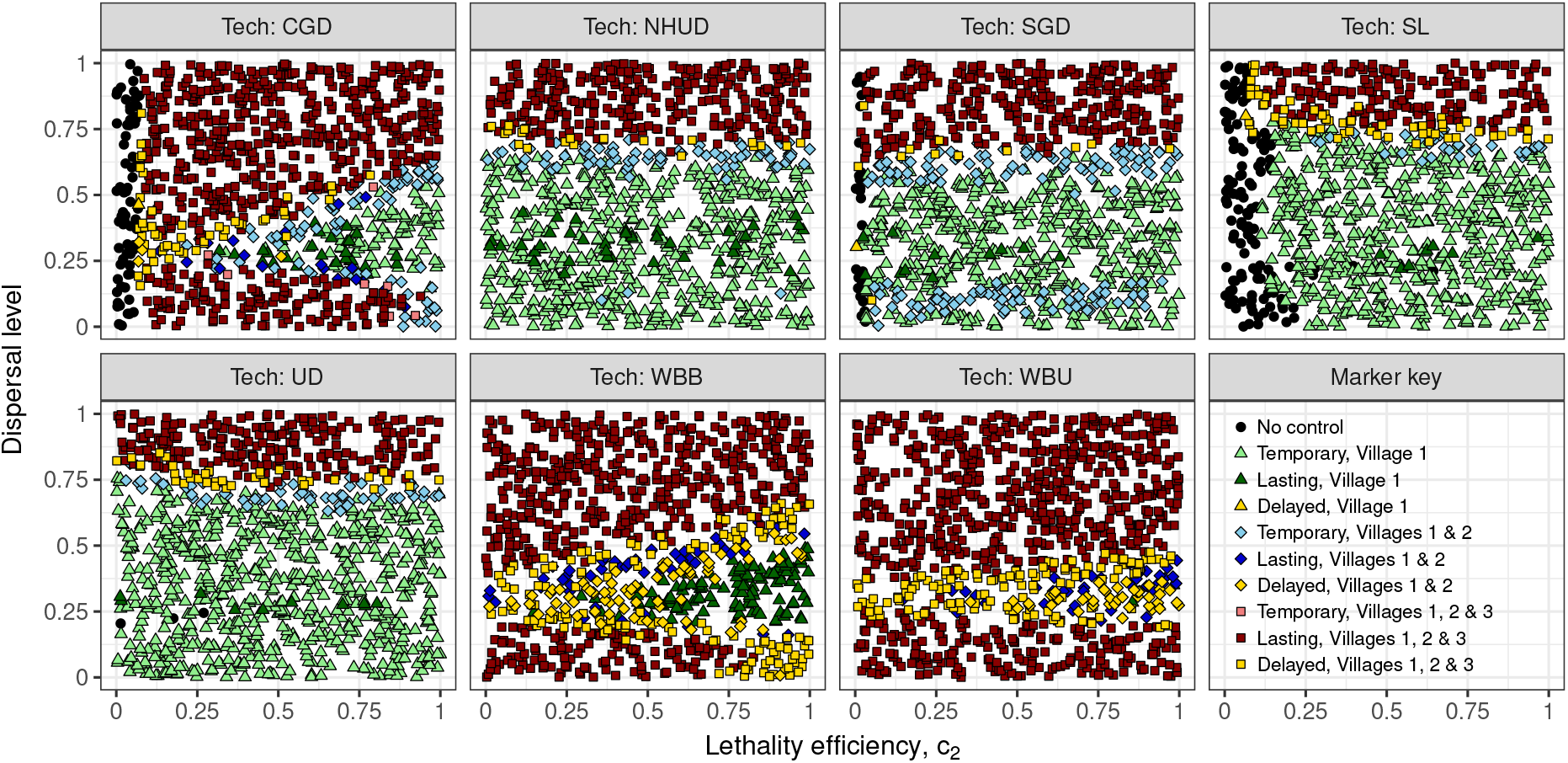
The outcomes of a year-long control effort in one village, with two neighbouring villages, in which 2000 modified mosquitoes were released every week, for varying values of the lethality efficiency, *c*_2_, of the payload genes and the dispersal level, which for values in [0, 1] scales the average percentage of mosquitoes migrating from each node each day between 0% and 15%, and scales the probability distribution governing the distances that migrants travel. The ambient fitness cost of a single transgene is *c*_1_ = 0.05. The mosquito simulated here is an ‘average’ species, with life history parameters taken as the mean of those listed for *Aedes* and *Anopheles* in table 1. The control technologies are a self-limiting technology (SL), homologous underdominance (UD), non-homologous underdominance (NHUD), a homing gene drive with combined homing and lethal gene (CGD), a homing gene drive with separated homing and lethal genes (SGD), unidirectional single-strain *Wolbachia* (WBU) and bidirectional two-strain *Wolbachia* (WBB). The bottom right panel shows marker shape and colour codings. Temporary control means control in the year of releases but not after releases stop; lasting control means control that continues after releases stop; delayed control means control that does not take affect until the year after releases stop.

To consider how variation in a species’ dispersal may affect whether and when a particular genetic control technology could be effective, we examine in the following results two mosquito archetypes with very different dispersal behaviours, which are broadly representative of the published dispersal ranges for *Aedes aegypti* and *Anopheles gambiae* [29]. For the “*Ae. aegypti* archetype” we assume that most mosquitoes fly less than 65m over their lifetime and ~ 1% fly over 400m [representative values from 64, 66]. For the “*An. gambiae* archetype”, we assume that most mosquitoes fly less than 800m and ~ 1% fly over 1.25km [representative values from 32, 34] (see appendix S1a for the mathematical details and fig. 15 for an example of average spatial spread over a single 30-day lifetime). The percentage of a population dispersing from their current node each day is randomly sampled for every node each day from a Gaussian distribution centred at 7.5% for both species archetypes.

### 3.2 Mating success

The assumption of random mating is inappropriate if released males have a lower mating success than wild males. We capture this possibility by allowing wild females to mate proportionally more with wild males than with any modified genotype (see appendix S3a), with decreasing ability to be choosy as the wild male population diminishes. Mosquito mating behaviour is poorly understood and therefore our model makes no assumptions about the number of mates a female has, male re-mating rates, male or female reproductive skew or fertilization rate arising from each copulation. We model variation in mating success as a change in the relative ability of transgenic males to sire viable offspring with wild females compared to the ability of wild males.

We find that mating success has a large impact on the success of genetic control. For all technologies except *Wolbachia*, a mating success lower than 30% prevents even local control of *An. gambiae* (fig. 3). The *Wolbachia* systems WBU and WBB are able to cause population replacement at lower values of mating success due to the release of modified females, allowing the infection to stay in the population even with low wild-type mating (due to the *Wolbachia*-infected population mating amongst themselves). Of note is the containing effect that a mating success of lower than ~ 70% has on the invasive CGD when the lethality efficiency is high; however, a moderate to low lethality efficiency of the payload gene acts to make the CGD invasive down to a mating success level of around 50% (fig. 4).

**Figure 3.**
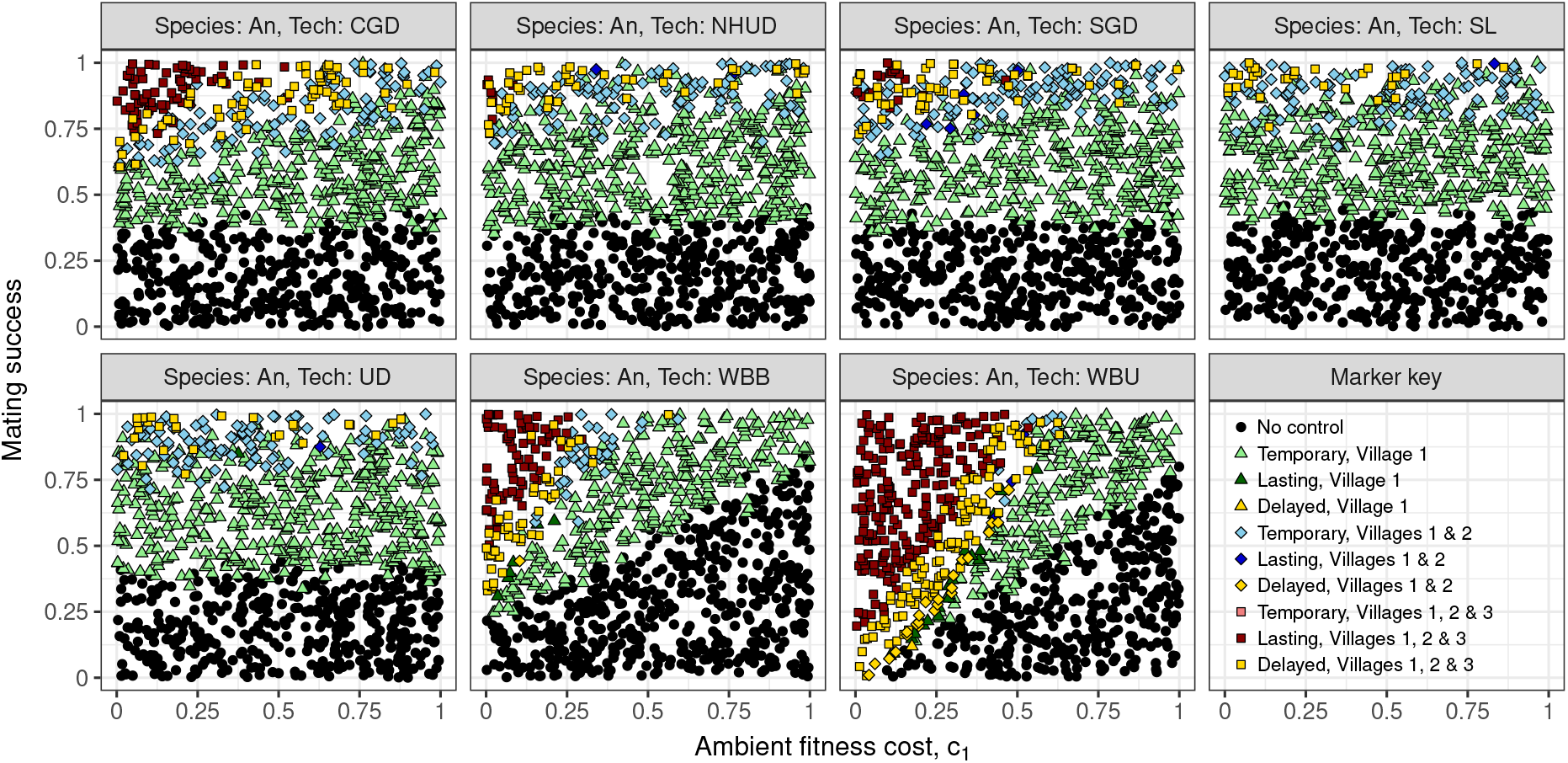
Outcomes of a year-long control effort for *An. gambiae* when the mating success of modified males and the ambient fitness cost, *c*_1_, imposed by each modified allele are varied. Mating success is defined in terms of the fraction of wild females who preferentially mate with wild-type males over males of a modified genotype: a mating success of one implies there is no mating preference; a mating success of zero implies all wild-type females choose to mate with wild-type males (see appendix S3a). Mating preference is scaled down when wild-type males are hard to find. The lethality efficiency of the payload genes is *c*_2_ = 0.85. The control technologies are a self-limiting technology (SL), homologous underdominance (UD), non-homologous underdominance (NHUD), a homing gene drive with combined homing and lethal gene (CGD), a homing gene drive with separated homing and lethal genes (SGD), unidirectional single-strain *Wolbachia* (WBU) and bidirectional two-strain *Wolbachia* (WBB). The bottom right panel shows marker shape and colour codings. Temporary control means control in the year of releases but not after releases stop; lasting control means control that continues after releases stop; delayed control means control that does not take affect until the year after releases stop.

**Figure 4.**
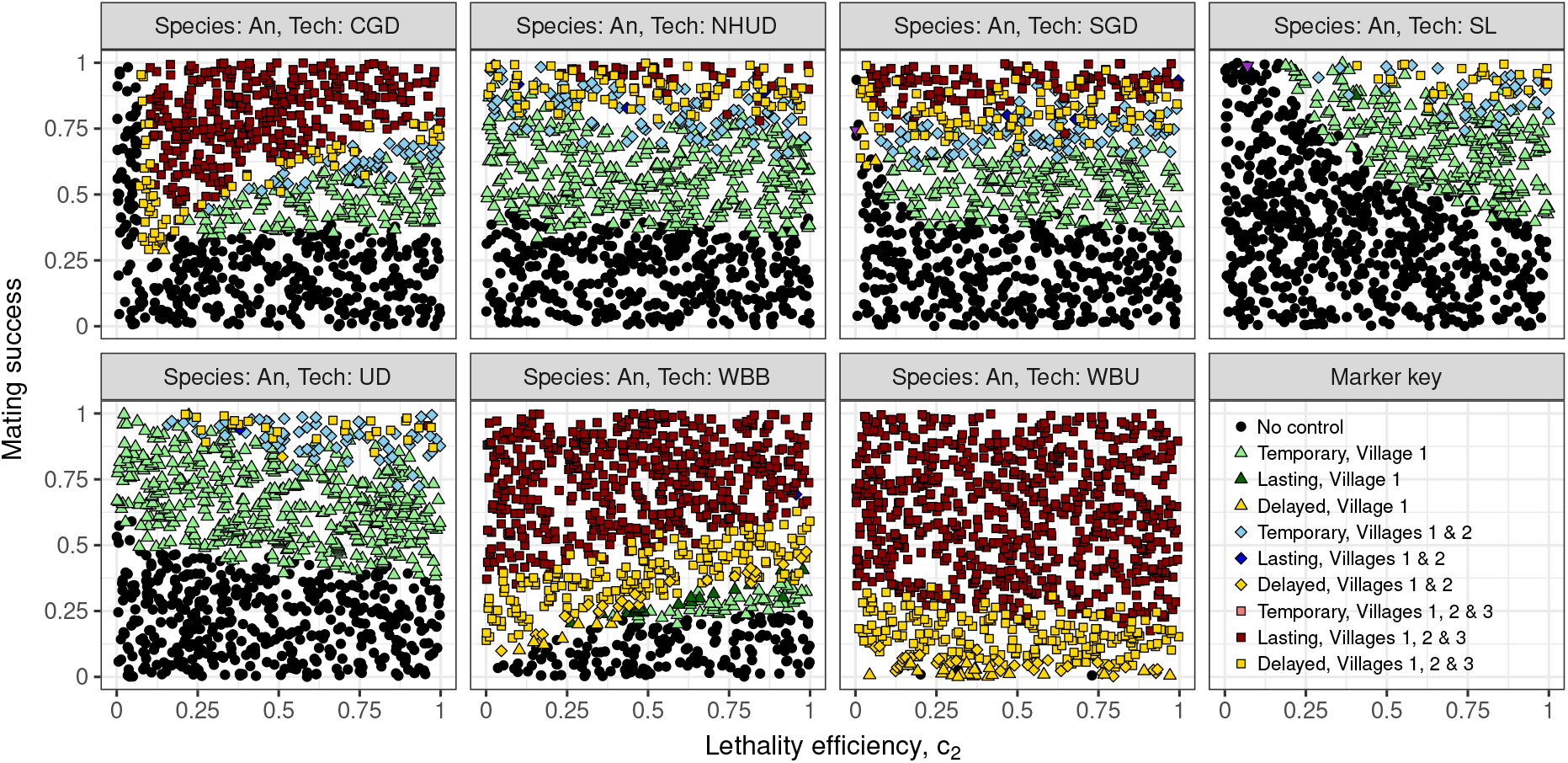
Outcomes of a year-long control effort for *An. gambiae* when the mating success of modified males and the lethality efficiency, *c*_2_, of the genetic control technologies are varied. Mating success is defined in terms of the fraction of wild females who preferentially mate with wild-type males over males of a modified genotype: a mating success of one implies there is no mating preference; a mating success of zero implies all wild-type females choose to mate with wild-type males (see appendix S3a). Mating preference is scaled down when wild-type males are hard to find. The ambient fitness cost of a single transgene is *c*_1_ = 0.05. The control technologies are a self-limiting technology (SL), homologous underdominance (UD), non-homologous underdominance (NHUD), a homing gene drive with combined homing and lethal gene (CGD), a homing gene drive with separated homing and lethal genes (SGD), unidirectional single-strain *Wolbachia* (WBU) and bidirectional two-strain *Wolbachia* (WBB). The bottom right panel shows marker shape and colour codings. Temporary control means control in the year of releases but not after releases stop; lasting control means control that continues after releases stop; delayed control means control that does not take affect until the year after releases stop.

These results also provide further evidence that spatial containment of any control technology may be easier in *Ae. aegypti* than *An. gambiae*. The SL, UD and NHUD achieve temporary, local suppression/replacement for a mating success above 40% in *Ae. aegypti* (figs. 5 and 6). For the CGD, global suppression of *Ae. aegypti* is achieved when the lethality efficiency is low and mating is almost random: for such a low lethality efficiency the gene drive is effectively acting as a weak bi-sex lethal technology, as the ambient fitness cost (which acts on males) is of the same order of magnitude (fig. 6). At very low mating success, the WBU is able to achieve population replacement while the the WBB is not. This is because of a population threshold effect: the releases under WBU are all of a single infection strain and the release sizes are enough to tip the balance in favour of the infection; in WBB the release size is split between two infection types, and the threshold effect does not trigger in favour of either infection strain. Widespread replacement across the three villages is achieved by both *Wolbachia* strategies in *Ae. aegypti* for near-random mating due to rare stochastic migration between villages (fig. 6).

**Figure 5.**
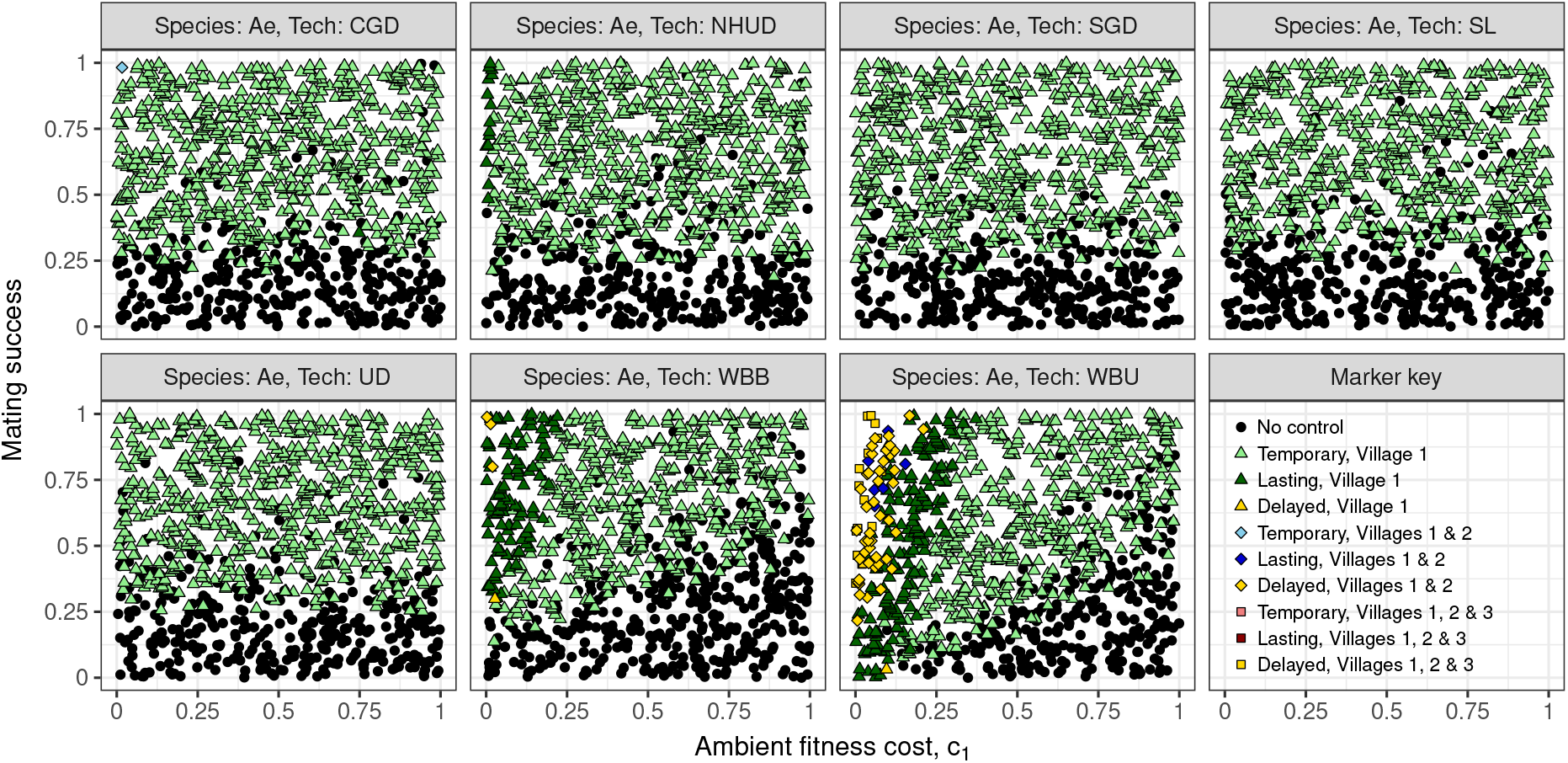
Outcomes of a year-long control effort for *Ae. aegypti* mosquitoes when the mating success of modified males and the ambient fitness cost, *c*_1_, imposed by each modified allele are varied. Mating success is defined in terms of the fraction of wild females who preferentially mate with wild-type males over males of a modified genotype: a mating success of one implies there is no mating preference; a mating success of zero implies all wild-type females choose to mate with wild-type males (see appendix S3a). Mating preference is scaled down when wild-type males are hard to find. The lethality efficiency of the payload genes is *c*_2_ = 0.85. The control technologies are a self-limiting technology (SL), homologous underdominance (UD), non-homologous underdominance (NHUD), a homing gene drive with combined homing and lethal gene (CGD), a homing gene drive with separated homing and lethal genes (SGD), unidirectional single-strain *Wolbachia* (WBU) and bidirectional two-strain *Wolbachia* (WBB). The bottom right panel shows marker shape and colour codings. Temporary control means control in the year of releases but not after releases stop; lasting control means control that continues after releases stop; delayed control means control that does not take affect until the year after releases stop.

**Figure 6.**
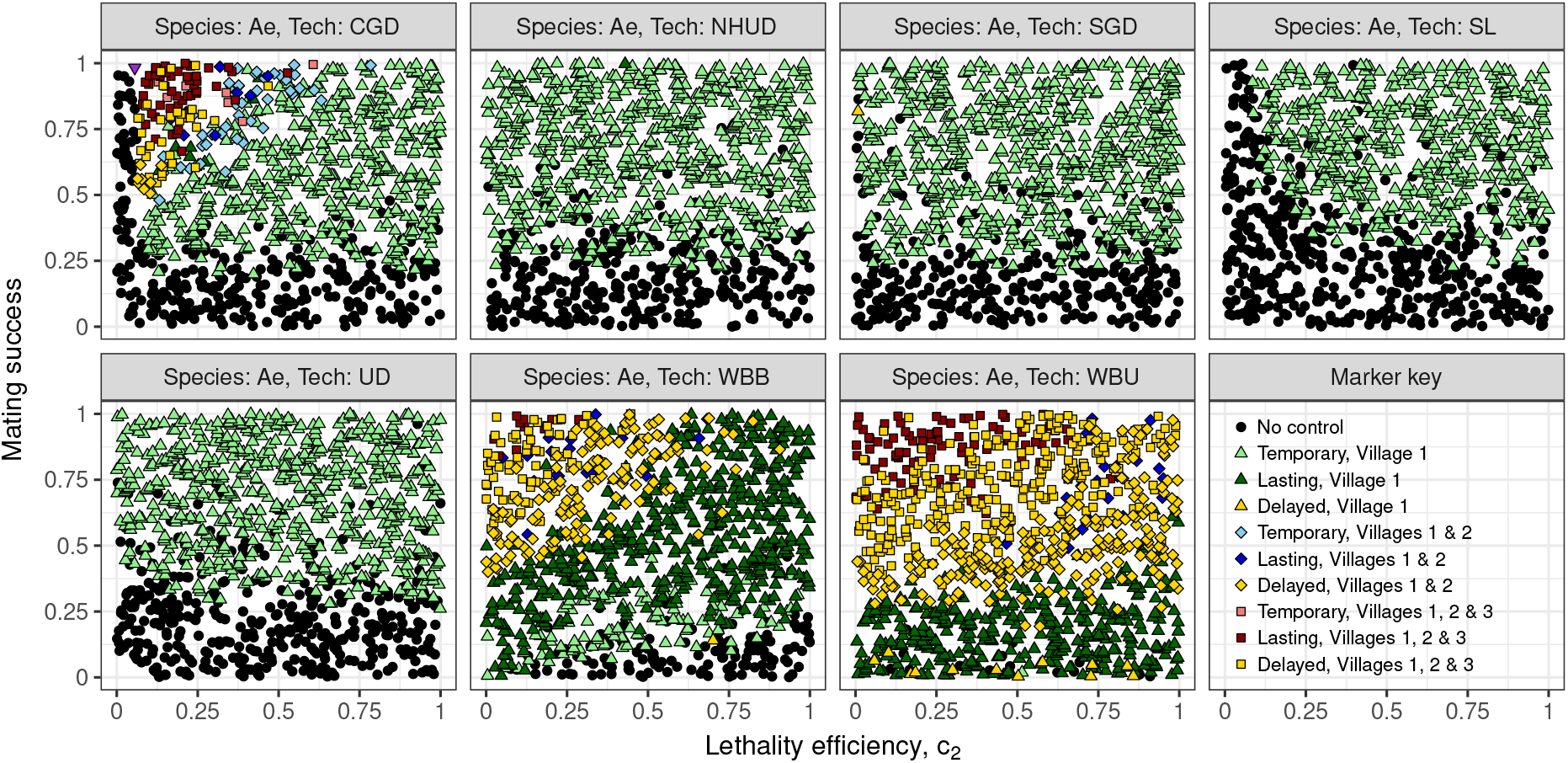
Outcomes of a year-long control effort for *Ae. aegypti* when the mating success of modified males and the lethality efficiency, *c*_2_, of the genetic control technologies are varied. Mating success is defined in terms of the fraction of wild females who preferentially mate with wild-type males over males of a modified genotype: a mating success of one implies there is no mating preference; a mating success of zero implies all wild-type females choose to mate with wild-type males (see appendix S3a). Mating preference is scaled down when wild-type males are hard to find. The ambient fitness cost of a single transgene is *c*_1_ = 0.05. The control technologies are a self-limiting technology (SL), homologous underdominance (UD), non-homologous underdominance (NHUD), a homing gene drive with combined homing and lethal gene (CGD), a homing gene drive with separated homing and lethal genes (SGD), unidirectional single-strain *Wolbachia* (WBU) and bidirectional two-strain *Wolbachia* (WBB). The bottom right panel shows marker shape and colour codings. Temporary control means control in the year of releases but not after releases stop; lasting control means control that continues after releases stop; delayed control means control that does not take affect until the year after releases stop.

### 3.3 *Anopheles* population size

Population genetic studies consistently show that the effective population size for *Anopheles* is ≳ 10000 [5] whereas for *Aedes* it is, on average, 400–600 [61]; their population census sizes are many tens of thousands and just a few thousand, respectively [67, 72]. Here we investigate how this discrepancy in population size affects the success of control efforts, by varying the base carrying capacity (prior to seasonality and environmental stochasticity) of all nodes. A steep gradient of greater than one separating the red and yellow suppression regions of the plots from the black regions means that an increase in release size has a larger effect than a relative increae in the wild population (release growth dominates) – this is true for WBU and CGD. A shallow gradient of less than one indicates that the growing population size swamps the relative growth in the size of releases (wild type growth dominates), such as for SL and UD (fig. 7).

**Figure 7.**
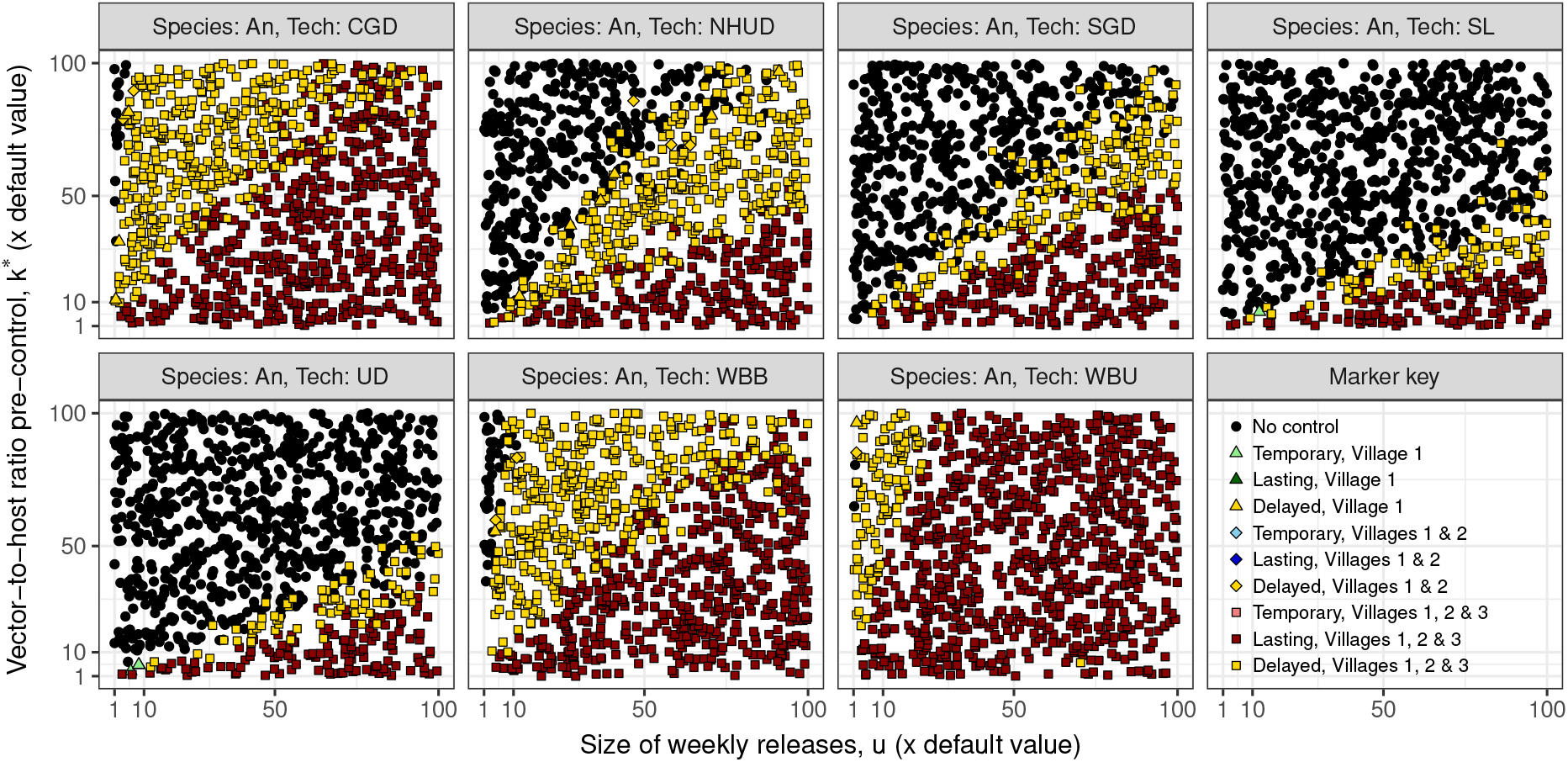
The effect on control efforts of varying the carrying capacity of each node, as weekly release sizes are varied. Scales are relative to the default parameter values in table 1. The ambient fitness cost of a single transgene is *c*_1_ = 0.05 and the lethality efficiency of the payload gene is *c*_2_ = 0.85. The mosquito species used is *An. gambiae*. The control technologies are a self-limiting technology (SL), homologous underdominance (UD), non-homologous underdominance (NHUD), a homing gene drive with combined homing and lethal gene (CGD), a homing gene drive with separated homing and lethal genes (SGD), unidirectional single-strain *Wolbachia* (WBU) and bidirectional two-strain *Wolbachia* (WBB). The bottom right panel shows marker shape and colour codings. Temporary control means control in the year of releases but not after releases stop; lasting control means control that continues after releases stop; delayed control means control that does not take affect until the year after releases stop.

### 3.4 Resistance and invasion

That mosquitoes will develop resistance to genetic control technologies (as they have developed resistance to insecticides and diminished the effecicacy of bed nets) is widely agreed upon. The speed with which this resistance will arise in the wild – either through the evolution of behavioural patterns that reduce the spread of modified genes, or through the chance generation of homing-resistance alleles through non-homologous end joining after DNA cutting [19, 68] – is unknown. Here we investigate how the speed with which behavioural resistance emerges (through the development of mate preference over time) affects the success of control efforts. We simulate over the parameter space spanned by the initial level of mating preference and the period of time over which the mating preference becomes total (100% of wild females choose to mate preferentially with wild males, unless lack of wild males forces them to mate with modified males).

The immediate development of strong mating preference causes a problem for all technologies and species (figs. 8 and 9). Only for very gradual emergence of behavioural resistance, in which it takes longer than two years to reach zero mating success (when there are sufficient wild males to mate with instead), can the CGD invade and suppress all three villages. Increasing release sizes is one way to combat this behavioural resistance, although the effect is not linear: diminishing returns can be seen when releasing larger numbers at low levels of mating success (figs. 10 and 11).

**Figure 8.**
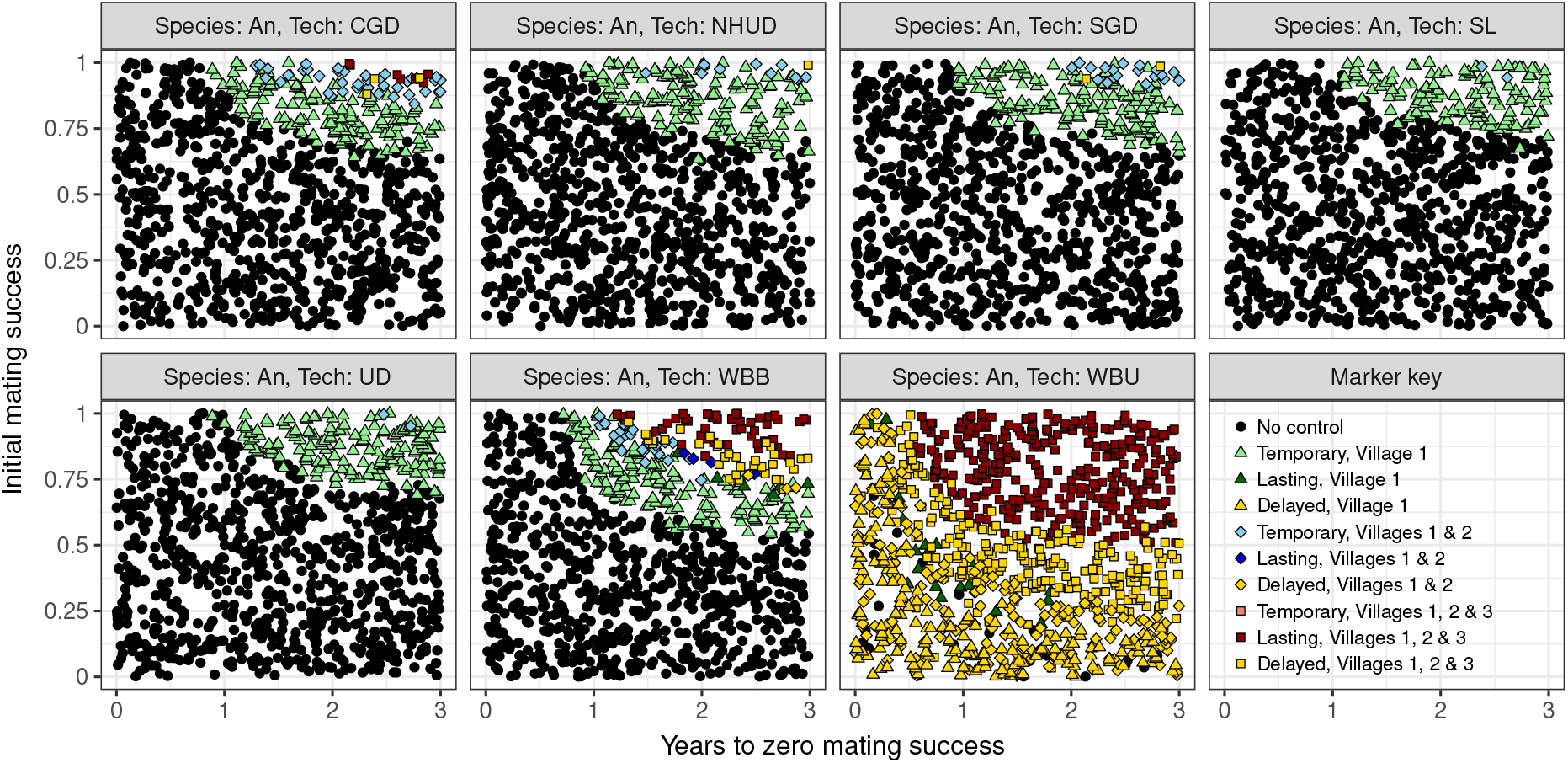
Varying the initial modified male mating success and the speed with which the mating success linearly decreases to zero (at which point wild females display complete behavioural resistance to the technologies), expressed in terms of the time taken for the rate to reach zero. The ambient fitness cost of a single transgene is *c*_1_ = 0.05 and the lethality efficiency of the payload gene is *c*_2_ = 0.85. The mosquito species used is *An. gambiae*. The control technologies are a self-limiting technology (SL), homologous underdominance (UD), non-homologous underdominance (NHUD), a homing gene drive with combined homing and lethal gene (CGD), a homing gene drive with separated homing and lethal genes (SGD), unidirectional single-strain *Wolbachia* (WBU) and bidirectional two-strain *Wolbachia* (WBB). The bottom right panel shows marker shape and colour codings. Temporary control means control in the year of releases but not after releases stop; lasting control means control that continues after releases stop; delayed control means control that does not take affect until the year after releases stop.

**Figure 9.**
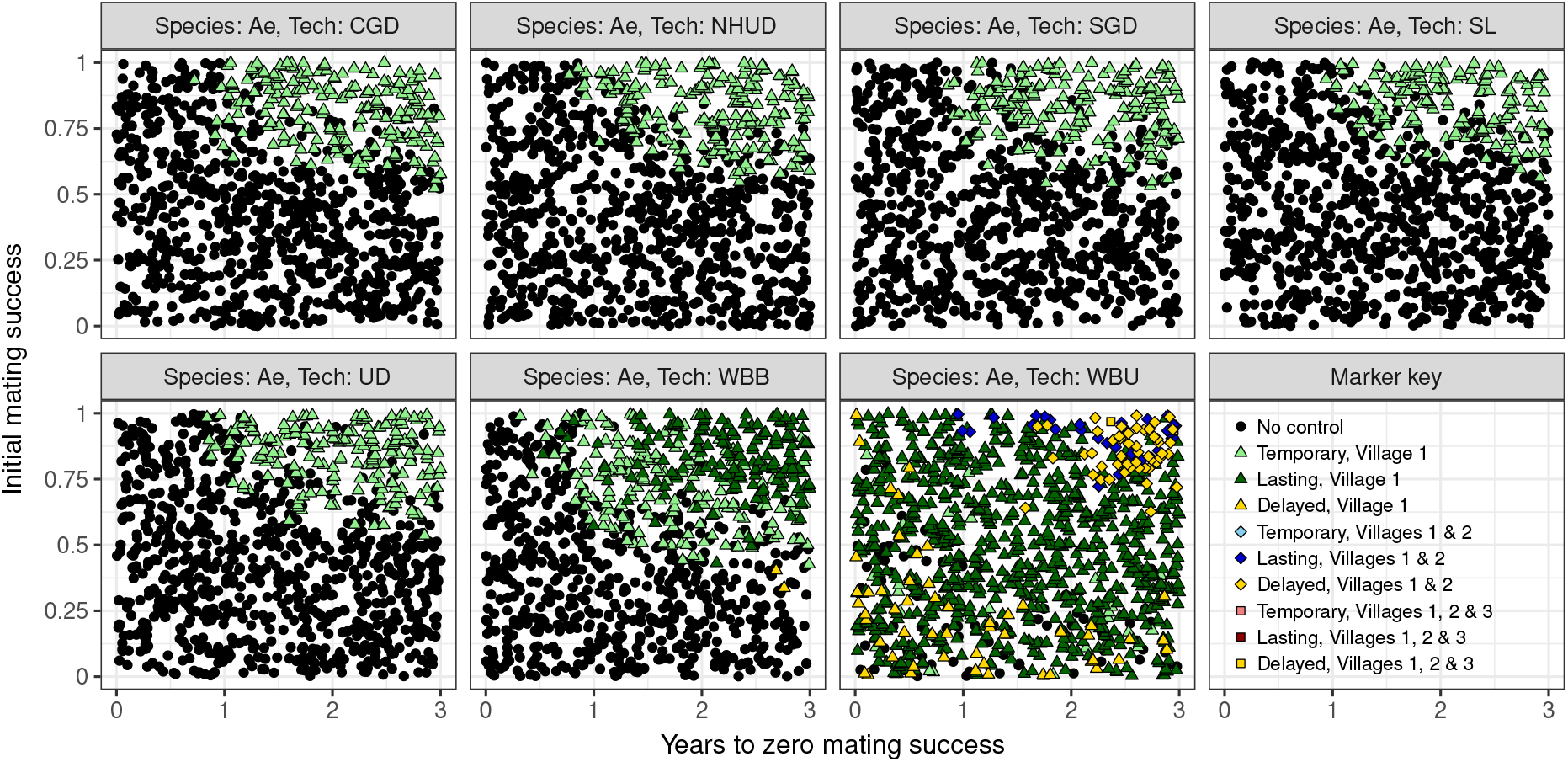
Varying the initial modified male mating success and the speed with which the mating success linearly decreases to zero (at which point wild females display complete behavioural resistance to the technologies), expressed in terms of the time taken for the rate to reach zero. The ambient fitness cost of a single transgene is *c*_1_ = 0.05 and the lethality efficiency of the payload gene is *c*_2_ = 0.85. The mosquito species used is *Ae. aegypti*. The control technologies are a self-limiting technology (SL), homologous underdominance (UD), non-homologous underdominance (NHUD), a homing gene drive with combined homing and lethal gene (CGD), a homing gene drive with separated homing and lethal genes (SGD), unidirectional single-strain *Wolbachia* (WBU) and bidirectional two-strain *Wolbachia* (WBB). The bottom right panel shows marker shape and colour codings. Temporary control means control in the year of releases but not after releases stop; lasting control means control that continues after releases stop; delayed control means control that does not take affect until the year after releases stop.

**Figure 10.**
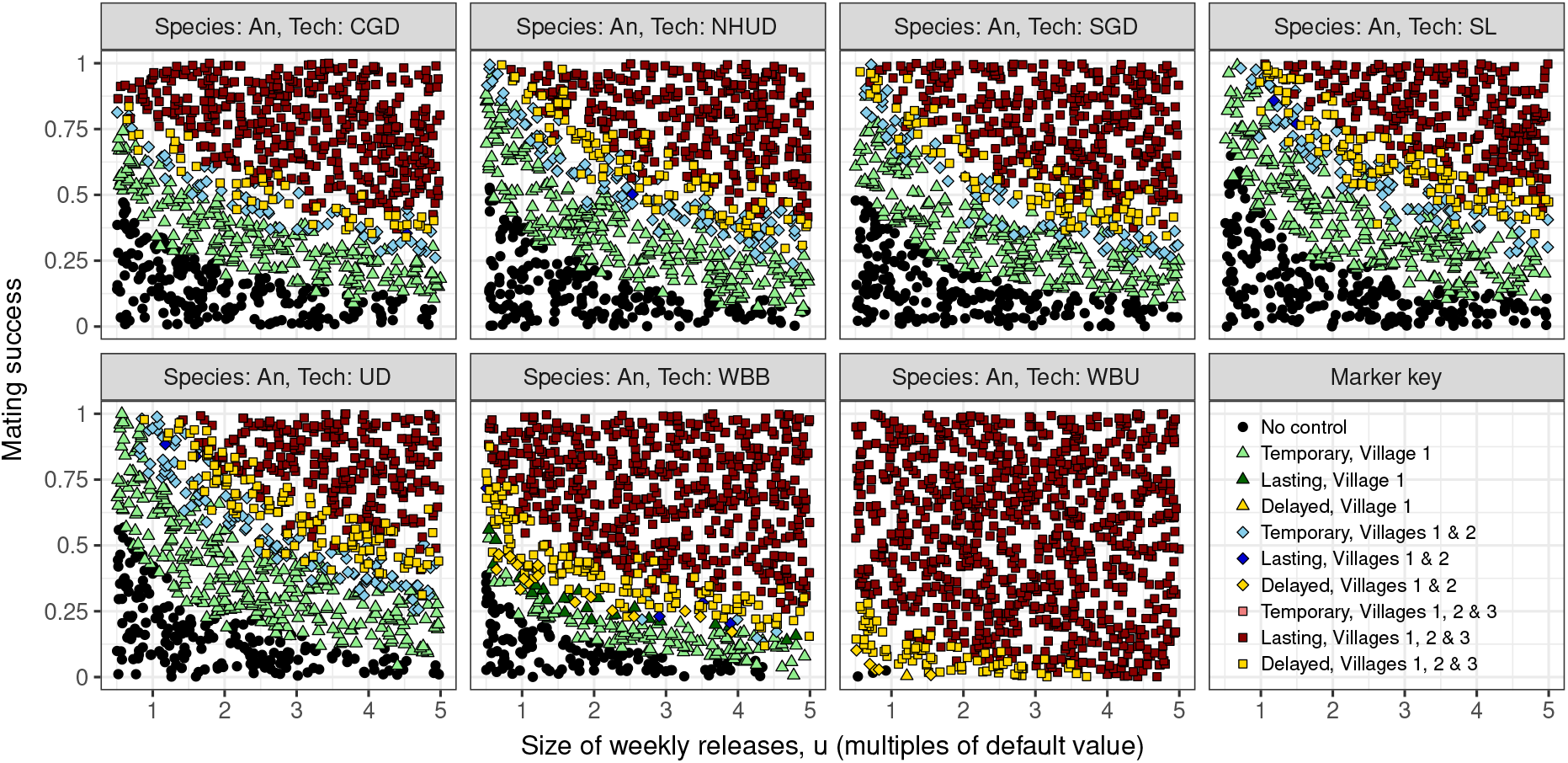
The effect of increasing release sizes, relative to the default value in table 1, to combat mating preference, for *An. gambiae*. The ambient fitness cost of a single transgene is *c*_1_ = 0.05 and the lethality efficiency of the payload gene is *c*_2_ = 0.85. The control technologies are a self-limiting technology (SL), homologous underdominance (UD), non-homologous underdominance (NHUD), a homing gene drive with combined homing and lethal gene (CGD), a homing gene drive with separated homing and lethal genes (SGD), unidirectional single-strain *Wolbachia* (WBU) and bidirectional two-strain *Wolbachia* (WBB). The bottom right panel shows marker shape and colour codings. Temporary control means control in the year of releases but not after releases stop; lasting control means control that continues after releases stop; delayed control means control that does not take affect until the year after releases stop.

**Figure 11.**
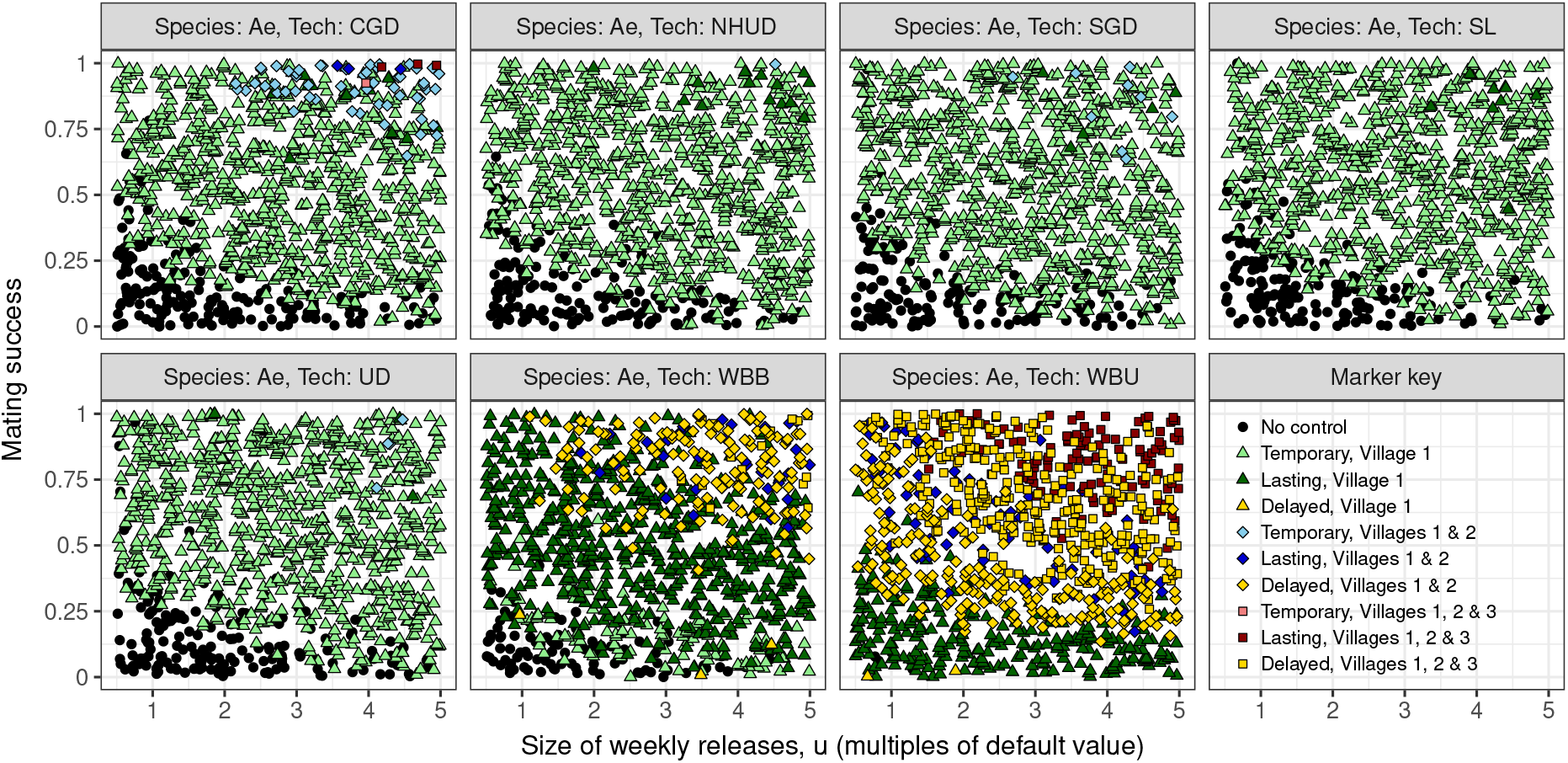
The effect of increasing release sizes, relative to the default value in table 1, to combat mating preference, for *Ae. aegypti*. The ambient fitness cost of a single transgene is *c*_1_ = 0.05 and the lethality efficiency of the payload gene is *c*_2_ = 0.85. The control technologies are a self-limiting technology (SL), homologous underdominance (UD), non-homologous underdominance (NHUD), a homing gene drive with combined homing and lethal gene (CGD), a homing gene drive with separated homing and lethal genes (SGD), unidirectional single-strain *Wolbachia* (WBU) and bidirectional two-strain *Wolbachia* (WBB). The bottom right panel shows marker shape and colour codings. Temporary control means control in the year of releases but not after releases stop; lasting control means control that continues after releases stop; delayed control means control that does not take affect until the year after releases stop.

*Wolbachia* is unique among the current genetic control technologies (not least because it is not a genetic control technology) due to the possibility of infection causing a fitness advantage in the modified mosquitoes over the wild type. This may be due to the bacterium causing the mosquito to be less vulnerable to viral infection [33, 65] and driving a higher larval survivorship in less competitive environments [27].

Here we study how the fitness differential between wild type and *Wolbachia*-infected mosquitoes affects the ability of the bacteria to invade neighbouring populations.

If there is close to zero infection leakage (the fraction of offspring that are wild type when they would normally have been infected if infection was completely efficient), even a small fitness advantage (shown as a negative fitness cost) for the *Wolbachia*-infected mosquitoes is very likely to lead to complete population replacement in the three villages (fig. 12). For *Ae. aegypti* mosquitoes, neutral or slightly costly *Wolbachia* strains coupled with a small amount of infection leakage is likely to lead to contained, and possibly temporary, population replacement. In all cases the unidirectional system is more invasive, even at relatively high fitness costs in the *An. gambiae* case. Inefficiencies in *Wolbachia* infection could prevent population replacement from taking place, even for very advantageous strains. If one is not able to alter the infection efficiency of *Wolbachia* strains, invasiveness could be managed by reducing release sizes. Containment can be achieved in the *Anopheles* case by halving the weekly release size to 1000 if the strains impose a moderate ambient fitness cost (fig. 13). However, even for the *Aedes* mosquitoes, *Wolbachia* strains with a moderate fitness advantage can invade neighbouring populations when releases are small.

**Figure 12.**
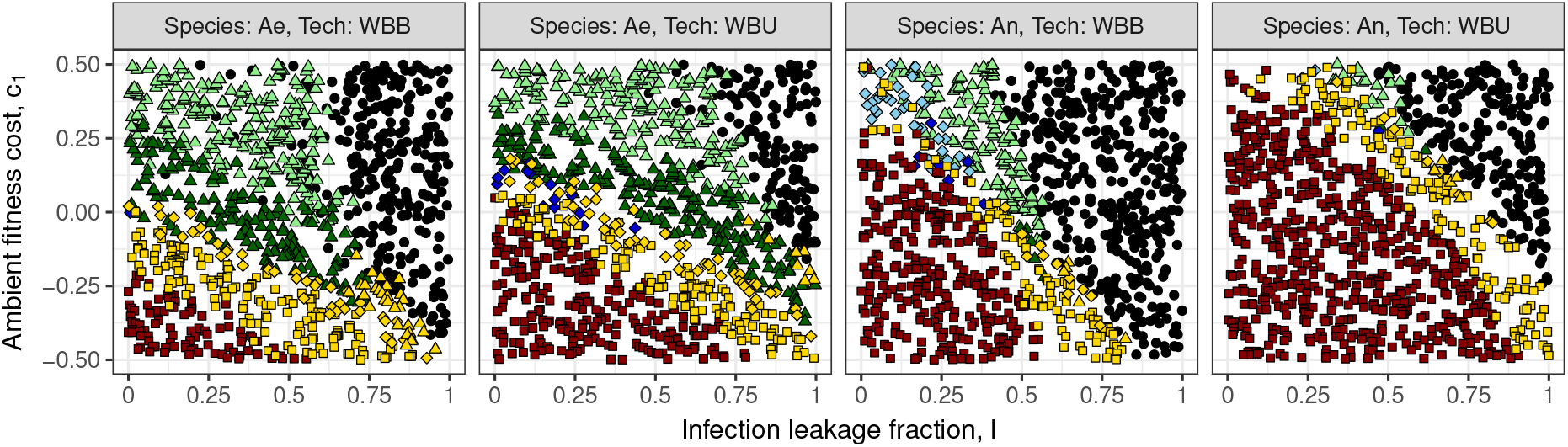
The effect of ambient fitness cost (*c*_1_ > 0) or advantage (*c*_1_ < 0) on the ability of two *Wolbachia* systems, unidirectional (WBU) and bidirectional (WBB), to invade neighbouring *Ae. aegypti* (Ae) and *An. gambiae* (An) populations, as the infection leakage of the *Wolbachia* strains is varied. We define infection leakage as the percentage of offspring that remain uninfected when they would usually be infected with *Wolbachia* (and can be thought of as the opposite of a homing rate). Marker colour and shape coding as in all other figures.

**Figure 13.**
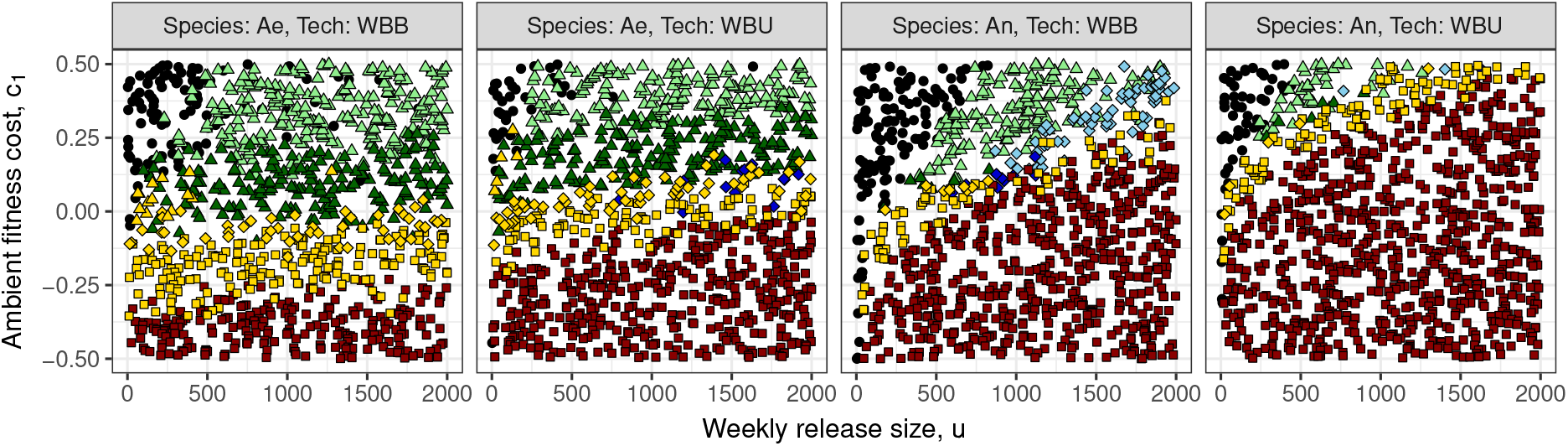
The effect of ambient fitness cost (*c*_1_ > 0) or advantage (*c*_1_ < 0) on the ability of two *Wolbachia* systems, unidirectional (WBU) and bidirectional (WBB), to invade neighbouring *Ae. aegypti* (Ae) and *An. gambiae* (An) populations, as the size of weekly releases of *Wolbachia*-infected males and females is varied. Releases are spead equally between both sexes in the unidirectional system and between both sexes and both strains in the bidirectional system. Marker colour and shape coding as in all other figures.

## 4 Discussion

We compared the ability of seven vector control technologies to suppress *Aedes aegypti* or *Anopheles gambiae* mosquito populations across three villages (or neighbourhoods), and examined the impact of mosquito dispersal behaviour and emergent behavioural resistance on the efficacy of genetic control.

We find that in mosquitoes with dispersal behaviour akin to *An. gambiae* (with an average lifetime dispersal in the range of several hundred metres to one kilometre), that it will be very difficult to contain a homing gene drive (in which the homing and lethal gene are combined) and *Wolbachia* systems if mating between modified and wild mosquitoes occurs successfully. This is a robust result: over a large range of population sizes and gene drive fitness costs, *Wolbachia* and a combined homing gene drive are ‘uncontained’ and will spread and cause lasting suppression across a large geographic area (fig. 1). Homing gene drives in which the drive gene and payload gene are separated (with the drive gene inherited at a Mendelian rate) are less invasive than combined gene drive systems. Spatial containment is likely with underdominance threshold drives and self-limiting technology although not guaranteed for highly dispersive species (fig. 2).

In contrast, we show that for mosquitoes with dispersal behaviour akin to *Ae. aegypti* (with an average lifetime dispersal of less than 100m), the invasiveness of a homing technology (and the invasive *Wolbachia* systems) is more dependent on the genetic fitness costs it imposes, and a complex balance between local suppression/replacement and repopulation from neighbouring wild-type populations exists (fig. 1). Note, we do not take human movement into account—possibly, a person could unintentionally introduce genetically modified mosquitoes into a distant control-naïve population. For *Ae. aegypti*, our results suggest that to achieve population suppression over a large geographic area will require multiple release locations and careful planning. This is in direct agreement with the findings of Legros et al. [47], in which large-area control of *Ae. aegypti* via female-killing transgenes (similar to the self-limiting technology studied here) was found to be an unrealistic goal unless spatially homogeneous releases could be performed. While this release protocol for *Ae. aegypti* may be more costly, the pay-off is that population suppression is likely to be sustained after releases stop due to the low level of immigration from uncontrolled areas—although long-term re-population is unavoidable [47]. In summary, these results suggests that there might be scope for safe deployment of so-called ‘global’ gene drives in one species, but not another. Put simply, a ‘one size fits all’ approach to genetic control will not work.

Together, our results demonstrate that the efficacy and safety of genetic control is extremely sensitive to biological and ecological details. For example, the stark differences in the impact of genetic control on *An. gambiae* and *Ae. aegypti* arise from how we assume these two species disperse and their different demographic parameters. At present, our understanding of mosquito dispersal is incomplete: within-species estimates of dispersal vary widely, the frequency of long-range dispersal in *An. gambiae* is unclear [15] and the impact of man-made obstacles and natural topography on movement is poorly understood. What is certain is that there is a significant difference in the dispersal behaviour of these species. Genetic studies show that the effective population size for *Ae. aegypti* populations is 300–600 and the census size is a few thousand [72], whereas in *An. gambiae* the effective population size is estimated to be ≳10,000 [5] and the census size is many tens of thousands [67]. Consequently, the speed and extent of gene flow through an *Ae. aegypti* population is vastly different to that in *An. gambiae*, and this may mean that different genetic control approaches and different spatial release strategies will be required. Our finding that mating success has a major impact on the efficacy of genetic control is particularly significant given that so little is known about mosquito mating behaviour. For example, we still do not know the degree to which females are able to choose their mates.

Our goal in this work was to demonstrate how the performance of different types of genetic control will be determined by the target species’ biology and ecology. In our mathematical approach we chose to make simplifying assumptions and generalizations about the life cycle and behaviour of mosquitoes. This is common in the literature, and we do it here in order to examine specific questions within a simplified framework that does not rely on the detailed, difficult and error-prone parameterisations of more complex agent-based models [46, 48, 56, 59]. Of course, it is vital that we understand the limitations of simplified models and that making common simplifying assumptions – while useful – should never obscure the importance of gaining a richer empirical understanding. First, we assume a simple life-cycle model with constant maturation rate and adult death rate; however, survivorship and maturation rate will be dependent on local conditions such as resource availability and temperature—which may affect wild and transgenic mosquitoes differently. A richer life-cycle model may be able to capture the phenotypic effect of genetic control more accurately, e.g. whether it reduces adult life spans, prevents successful oviposition or increases larval death rates, and hence could elucidate how changes in the local ecology affect their action. We captured seasonal changes in population size by varying the carrying capacity of each node; however seasonality could impact the life-cycle in many ways: changing temperature and the resulting effects on maturation; the number and quality of breeding sites available [as impressively modelled in current agent-based approaches 48, 56]; the intensity of predation; and the availability of sugar. In our model we assume that the host population remains constant in time and equal across all spatial sites. However, seasonal human migration into and out of disease hotspots is common – due to, for instance, pastoralism – and the interaction between human movement and mosquito seasonal dynamics may pose a unique challenge for disease control programmes [57, 58].

Second, we assume random, continuous mating and capture variation in mating success between wild types and transgenics simply as a shift in offspring proportions. This variation in mating success could arise from differences in re-mating behaviour, disparity in fertilization rates, numbers of viable progeny or female mate choice. A theoretical and empirical understanding of these biological details is needed. For example, if variation in mating success is mainly due to numbers of viable progeny, the impact on genetic control strategies may be limited to adjusting release rates. However, if female mate choice is driving the variation in mating success, this may have important consequences for long-term genetic control. If wild females can avoid or reject transgenic males then natural selection on female choice could cause genetic control to fail due to ‘behavioural resistance’ (which would manifest in the current model as a significant decrease in the mating success of released mosquitoes). Females have been shown to exhibit clear rejection behaviours towards males. In species such as *Ae. aegypti*, where males do not provide material resources such as food or territory, females are predicted to assess and choose mates based on heritable traits associated with improved offspring fitness [11, 42]. Our results suggest that even small reductions in the mating success of genetically modified males with wild females – due to a rapid appearance of behavioural resistance, for example – may prevent the spread of homing gene drives, creating a spatial containment effect (figs. 3–4 and 6). If this behavioural resistance emerges very quickly (within a year or two of the start of releases), it could damage the efficacy of control efforts to a large degree (figs. 8 and 9), and this damage can only partially be prevented by increasing release sizes (figs. 10 and 11).

Third, we assume unintended fitness costs are constant in time. However, at least for some control technologies we would expect that these costs would increase over time. For example, this may occur in the case of a CRISPR/Cas9 homing gene drive, where unintended DNA mutations (off-target effects) build up over time and may lead to an increasing unintended fitness cost on gene drive-bearing insects. If these ambient fitness costs can be predicted on a generation-by-generation basis (or at a more granular resolution), computational models like that presented here could be used to inform how control strategies might be altered to account for the changing fitness burden.

Overall, these results show that a clearer understanding of the biology and ecology of target mosquito species will be vital for the successful implementation of genetic control for population suppression. The “success” of an implementation is measured here in terms of the scale, spatial extent and permanence/transience of vector population suppression (or replacement). In the field, these metrics of success could be assessed through mark-release-recapture operations, and the short-term efficacy of a control programme could be monitored. Over the longer term, governments and national and international health agencies will measure success through reductions in disease levels; easing of the burden on local health services; the lowering of the number of work/school days lost to illness; and the value for money that the control programme represents to achieve these improvements. Weighing costs and benefits of disease control schemes is vitally important [3]; finding optimal implementation strategies could minimise the economic burden while maintaining effective disease reduction [41]. However, variability in vector behaviour, environmental stochasticity and the possibility of disease reintroduction via human or mosquito migration mean there cannot be a single catch-all strategy for cost-effective vector control. Our results suggest that some forms of genetic control may be safe and effective in one species but not in another. Even within a species, the most suitable genetic control strategy will depend upon the local environment and ecology. This presents a challenge for regulators; any given genetic control approach may be effective and/or safely contained in one setting, but not in another, suggesting that a flexible application-specific regulatory approach is needed.

Risk assessment of GM mosquitoes is guided by national and international legislative instruments, implementation frameworks and guidance policies [14, 23, 37, 71]. Current assessment guidance, associated with only risks of GM mosquito releases, follows three main principles: (i) efficacy of the technology, (ii) biosafety associated with adverse effects of releases on human health and/or wider biodiversity and (iii) cultural acceptance of novel technologies [69]. Current assessment of GM mosquitoes by national and international public health agencies is under way [e.g., 70] and our work is entirely pertinent to these assessments as the spatio-temporal dynamics of different mosquitoes will feed into efficacy assessments (identifying key parameters that affect the outcome of mosquito control) and biosafety (understanding the implication of the wider impacts of GM mosquito releases on the public health burden of disease and the ecological effects of control on wider biodiversity). Given all this, we argue that there is now compelling scope to consider public health benefits together with the risks of modified mosquito releases within inclusive, proportionate legislative and guidance frameworks.

Optimistically, our comparative approach for assessing different mosquito control interventions provides a tool to adopt and develop for assessing risks and biosafety of different genetics-based approaches for integrated vector control and management (particularly in cases where the specific and detailed data required for parameterisation of more complex agent-based models are unavailable, which is commonly the case). To date, most theoretical studies have focused on finding the specifications of the biotechnology that produce the desired effects in simplified population genetics models or through complex individual-based models. By comparing across technologies and species, our results show that these specifications are not as important as the understudied, poorly understood ecological, evolutionary and environmental factors in determining whether genetic control will fail or succeed.

## Acknowledgments

We thank Detlef Bartsch, Laura Harrington and the Mathematical Ecology Research Group in Oxford for discussion and comments on this work. The work was supported by funding from DARPA (HR0011-16-2-0006)

## Conflict of interest

The authors declare that they have no conflict of interest.

## Notes

### Competing Interest Statement

The authors have declared no competing interest.

## References

[1] Alphey, L. (2002). Re-engineering the sterile insect technique. Insect Biochemistry and Molecular Biology, 32(10):1243–1247.

[2] Alphey, L. (2014). Genetic control of mosquitoes. Annual Review of Entomology, 59:205–224.

[3] Alphey, N., Alphey, L., and Bonsall, M. B. (2011). A model framework to estimate impact and cost of genetics-based sterile insect methods for dengue vector control. PLOS ONE, 6(10):1–12.

[4] Alphey, N. and Bonsall, M. B. (2014). Interplay of population genetics and dynamics in the genetic control of mosquitoes. Journal of The Royal Society Interface, 11(93).

[5] Athrey, G., Hodges, T. K., Reddy, M. R., Overgaard, H. J., Matias, A., Ridl, F. C., Kleinschmidt, I., Caccone, A., and Slotman, M. A. (2012). The effective population size of malaria mosquitoes: Large impact of vector control. PLOS Genetics, 8(12):1–14.

[6] Barreaux, A. M. G., Stone, C. M., Barreaux, P., and Koella, J. C. (2018). The relationship between size and longevity of the malaria vector *Anopheles gambiae* (s.s.) depends on the larval environment. Parasites & Vectors, 11(1):485.

[7] Bull, J. J. (2015). Evolutionary decay and the prospects for long-term disease intervention using engineered insect vectors. Evolution, Medicine, and Public Health, 2015(1):152–166.

[8] Burt, A. (2003). Site-specific selfish genes as tools for the control and genetic engineering of natural populations. Proceedings of the Royal Society B: Biological Sciences, 270(1518):921–928.

[9] Bushland, R. C. (1974). Letter: Screwworm eradication program. Science, 184(4140):1010–1011.

[10] Carvalho, D. O., McKemey, A. R., Garziera, L., Lacroix, R., Donnelly, C. A., Alphey, L., Malavasi, A., and Capurro, M. L. (2015). Suppression of a field population of aedes aegypti in brazil by sustained release of transgenic male mosquitoes. PLOS Neglected Tropical Diseases, 9(7):1–15.

[11] Cator, L. J. and Harrington, L. C. (2011). The Harmonic Convergence of Fathers Predicts the Mating Success of Sons in Aedes aegypti. Animal Behaviour, 82(4):627–633.

[12] Cho, S. W., Kim, S., Kim, Y., Kweon, J., Kim, H. S., Bae, S., and Kim, J. S. (2014). Analysis of off-target effects of CRISPR/Cas-derived RNA-guided endonucleases and nickases. Genome Research, 24(1):132–141.

[13] Clements, A. N. and Paterson, G. D. (1981). The analysis of mortality and survival rates in wild populations of mosquitoes. Journal of Applied Ecology, 18(2):373–399.

[14] Council of European Union (2001). Council Directive 2001/18/EC on the deliberate release into the environment of genetically modified organisms. OJ L 106/0001 - 0039.

[15] Dao, A., Yaro, A. S., Diallo, M., Timbine, S., Huestis, D. L., Kassogue, Y., Traore, A. I., Sanogo, Z. L., Samake, D., and Lehmann, T. (2014). Signatures of aestivation and migration in Sahelian malaria mosquito populations. Nature, 516(7531):387–390.

[16] Davis, S., Bax, N., and Grewe, P. (2001). Engineered underdominance allows efficient and economical introgression of traits into pest populations. Journal of Theoretical Biology, 212(1):83–98.

[17] Davis, W. A., Clarke, P. M., Siba, P. M., Karunajeewa, H. A., Davy, C., Mueller, I., and Davis, T. M. E. (2011). Cost–effectiveness of artemisinin combination therapy for uncomplicated malaria in children: data from papua new guinea. Bulletin of the World Health Organization, 89(3):211–220.

[18] Deredec, A., Godfray, H. C. J., and Burt, A. (2011). Requirements for effective malaria control with homing endonuclease genes. Proceedings of the National Academy of Sciences, 108(43):E874–E880.

[19] Drury, D. W., Dapper, A. L., Siniard, D. J., Zentner, G. E., and Wade, M. J. (2017). CRISPR/Cas9 gene drives in genetically variable and nonrandomly mating wild populations. Science Advances, 3(5):e1601910.

[20] Dye, C. (1984). Models for the population dynamics of the yellow fever mosquito, aedes aegypti. Journal of Animal Ecology, 53(1):247–268.

[21] E., M. L. and H., K. B. (1998). *Aedes aegypti* survival and dispersal estimated by mark-release-recapture in northern australia. The American Journal of Tropical Medicine and Hygiene, 58(3):277–282.

[22] Eckhoff, P. A., Wenger, E. A., Godfray, H. C. J., and Burt, A. (2017). Impact of mosquito gene drive on malaria elimination in a computational model with explicit spatial and temporal dynamics. Proceedings of the National Academy of Sciences, 114(2):E255–E264.

[23] EFSA (2013). Guidance on the environmental risk assessment of genetically modified animals. EFSA Journal, 11(5):3200–3390.

[24] Ermert, V., Fink, A. H., Jones, A. E., and Morse, A. P. (2011a). Development of a new version of the liverpool malaria model. i. refining the parameter settings and mathematical formulation of basic processes based on a literature review. Malaria Journal, 10(1):35.

[25] Ermert, V., Fink, A. H., Jones, A. E., and Morse, A. P. (2011b). Development of a new version of the Liverpool Malaria Model. II. Calibration and validation for West Africa. Malaria Journal, 10(1):62.

[26] Esvelt, K. M., Smidler, A. L., Catteruccia, F., and Church, G. M. (2014). Emerging technology: Concerning rna-guided gene drives for the alteration of wild populations. eLife, 3:e03401.

[27] Gavotte, L., Mercer, D. R., Stoeckle, J. J., and Dobson, S. L. (2010). Costs and benefits of Wolbachia infection in immature Aedes albopictus depend upon sex and competition level. Journal of Invertebrate Pathology, 105(3):341–346.

[28] Gorman, K., Young, J., Pineda, L., Márquez, R., Sosa, N., Bernal, D., Torres, R., Soto, Y., Lacroix, R., Naish, N., Kaiser, P., Tepedino, K., Philips, G., Kosmann, C., and Cáceres, L. (2016). Short-term suppression of aedes aegypti using genetic control does not facilitate aedes albopictus. Pest Management Science, 72(3):618–628.

[29] Guerra, C. A., Reiner, R. C., Perkins, T. A., Lindsay, S. W., Midega, J. T., Brady, O. J., Barker, C. M., Reisen, W. K., Harrington, L. C., Takken, W., Kitron, U., Lloyd, A. L., Hay, S. I., Scott, T. W., and Smith, D. L. (2014). A global assembly of adult female mosquito mark-release-recapture data to inform the control of mosquito-borne pathogens. Parasites & Vectors, 7(1):276.

[30] Hancock, P. A., White, V. L., Callahan, A. G., Godfray, C. H. J., Hoffmann, A. A., and Ritchie, S. A. (2016a). Density-dependent population dynamics in aedes aegypti slow the spread of wmel wolbachia. Journal of Applied Ecology, 53(3):785–793.

[31] Hancock, P. A., White, V. L., Ritchie, S. A., Hoffmann, A. A., and Godfray, H. C. J. (2016b). Predicting wolbachia invasion dynamics in aedes aegypti populations using models of density-dependent demographic traits. BMC Biology, 14(1):96.

[32] Harrington, L. C., Scott, T. W., Lerdthusnee, K., Coleman, R. C., Costero, A., Clark, G. G., Jones, J. J., Kitthawee, S., Kittayapong, P., Sithiprasasna, R., and Edman, J. D. (2005). Dispersal of the dengue vector Aedes aegypti within and between rural communities. The American Journal of Tropical Medicine and Hygiene, 72(2):209–220.

[33] Hedges, L. M., Brownlie, J. C., O’Neill, S. L., and Johnson, K. N. (2008). Wolbachia and virus protection in insects. Science, 322(5902):702.

[34] Hemme, R. R., Thomas, C. L., Chadee, D. D., and Severson, D. W. (2010). Influence of urban landscapes on population dynamics in a short-distance migrant mosquito: evidence for the dengue vector Aedes aegypti. PLOS Neglected Tropical Diseases, 4(3):e634.

[35] Hibino, Y. and Iwahashi, O. (1991). Appearance of wild females unreceptive to sterilized males on Okinawa Is. in the eradication program of the melon fly, dacus cucurbitae COQUILLETT (Diptera: Tephritidae). Applied Entomology and Zoology, 26(2):265–270.

[36] Hoshen, M. B. and Morse, A. P. (2004). A weather-driven model of malaria transmission. Malaria Journal, 3(1):32.

[37] House of Lords Science & Technology Select Committee (2015). Genetically modified insects. Published by the Authority of the House of Lords, HL Paper 68, pp56.

[38] Huang, Y., Lloyd, A. L., Legros, M., and Gould, F. (2011). Gene-drive into insect populations with age and spatial structure: a theoretical assessment. Evolutionary Applications, 4(3):415–428.

[39] Jannat, K. N.-E. and Roitberg, B. D. (2013). Effects of larval density and feeding rates on larval life history traits in *Anopheles gambiae* s.s. (diptera: Culicidae). Journal of Vector Ecology, 38(1):120–126.

[40] Khamis, D., El Mouden, C., Kura, K., and Bonsall, M. B. (2018a). Ecological effects on underdominance threshold drives for vector control. Journal of Theoretical Biology, 456:1–15.

[41] Khamis, D., El Mouden, C., Kura, K., and Bonsall, M. B. (2018b). Optimal control of malaria: combining vector interventions and drug therapies. Malaria Journal, 17(1):174.

[42] Kokko, H., Brooks, R., Jennions, M. D., and Morley, J. (2003). The evolution of mate choice and mating biases. Proceedings of the Royal Society B: Biological Sciences, 270(1515):653–664.

[43] Kweka, E. J., Tenu, F., Magogo, F., and Mboera, L. E. G. (2015). Anopheles gambiae sensu stricto Aquatic Stages Development Comparison between Insectary and Semifield Structure. Advances in Zoology, 2015(720365):6.

[44] Lana, R. M., Carneiro, T. G. S., Honório, N. A., and Codeço, C. T. (2014). Seasonal and nonseasonal dynamics of aedes aegypti in rio de janeiro, brazil: Fitting mathematical models to trap data. Acta Tropica, 129:25–32. Human Infectious Diseases and Environmental Changes.

[45] Lee, H. L. and Joko, H. (2009). Comparative life parameters of transgenic and wild strain of *Aedes aegypti* in the laboratory. WHO Regional Office for South-East Asia.

[46] Legros, M., Xu, C., Morrison, A., Scott, T. W., Lloyd, A. L., and Gould, F. (2013). Modeling the dynamics of a non-limited and a self-limited gene drive system in structured aedes aegypti populations. PLOS ONE, 8(12).

[47] Legros, M., Xu, C., Okamoto, K., Scott, T. W., Morrison, A. C., Lloyd, A. L., and Gould, F. (2012). Assessing the feasibility of controlling Aedes aegypti with transgenic methods: a model-based evaluation. PLOS ONE, 7(12):e52235.

[48] Magori, K., Legros, M., Puente, M. E., Focks, D. A., Scott, T. W., Lloyd, A. L., and Gould, F. (2009). Skeeter Buster: a stochastic, spatially explicit modeling tool for studying Aedes aegypti population replacement and population suppression strategies. PLOS Neglected Tropical Diseases, 3(9):e508.

[49] Marshall, J. M. (2009). The effect of gene drive on containment of transgenic mosquitoes. Journal of Theoretical Biology, 258(2):250–265.

[50] Marshall, J. M. and Hay, B. A. (2012). Confinement of gene drive systems to local populations: a comparative analysis. Journal of Theoretical Biology, 294:153–171.

[51] Maynard Smith, J. and Slatkin, M. (1973). The stability of predator-prey systems. Ecology, 54(2):384–391.

[52] McInnis, D. O., Lance, D. R., and Jackson, C. G. (1996). Behavioral resistance to the sterile insect technique by mediterranean fruit fly (Diptera: Tephritidae) in Hawaii. Annals of the Entomological Society of America, 89:739–744.

[53] Midega, J. T., Mbogo, C. M., Mwnambi, H., Wilson, M. D., Ojwang, G., Mwangangi, J. M., Nzovu, J. G., Githure, J. I., Yan, G., and Beier, J. C. (2007). Estimating dispersal and survival of Anopheles gambiae and Anopheles funestus along the Kenyan coast by using mark-release-recapture methods. Journal of Medical Entomology, 44(6):923–929.

[54] Molineaux, L. and Gramiccia, G. (1980). The Garki project : research on the epidemiology and control of malaria in the Sudan savanna of West Africa. World Health Organization, Geneva.

[55] National Academies of Sciences, E. and Medicine (2016). Gene Drives on the Horizon: Advancing Science, Navigating Uncertainty, and Aligning Research with Public Values. The National Academies Press, Washington, DC.

[56] North, A., Burt, A., and Godfray, H. C. J. (2013). Modelling the spatial spread of a homing endonuclease gene in a mosquito population. Journal of Applied Ecology, 50(5):1216–1225.

[57] North, A. and Godfray, H. C. J. (2017). The dynamics of disease in a metapopulation: The role of dispersal range. Journal of Theoretical Biology, 418:57–65.

[58] North, A. R., Burt, A., and Godfray, H. C. J. (2019). Modelling the potential of genetic control of malaria mosquitoes at national scale. BMC Biology, 17(1):26.

[59] Oluwagbemi, O. O., Fornadel, C. M., Adebiyi, E. F., Norris, D. E., and Rasgon, J. L. (2013). ANOSPEX: a stochastic, spatially explicit model for studying Anopheles metapopulation dynamics. PLOS ONE, 8(7):e68040.

[60] O’Neill, S. L., Ryan, P. A., Turley, A. P., Wilson, G., Retzki, K., Iturbe-Ormaetxe, I., Dong, Y., Kenny, N., Paton, C. J., Ritchie, S. A., Brown-Kenyon, J., Stanford, D., Wittmeier, N., Anders, K. L., and Simmons, C. P. (2018). Scaled deployment of Wolbachia to protect the community from dengue and other Aedes transmitted arboviruses. Gates Open Res, 2:36.

[61] Saarman, N. P., Gloria-Soria, A., Anderson, E. C., Evans, B. R., Pless, E., Cosme, L. V., Gonzalez-Acosta, C., Kamgang, B., Wesson, D. M., and Powell, J. R. (2017). Effective population sizes of a major vector of human diseases, aedes aegypti. Evolutionary Applications, 10(10):1031–1039.

[62] Sheppard, P. M., Macdonald, W. W., Tonn, R. J., and Grab, B. (1969). The dynamics of an adult population of *Aedes aegypti* in relation to dengue haemorrhagic fever in bangkok. Journal of Animal Ecology, 38(3):661–702.

[63] Southwood, T. R. E., Murdie, G., Yasuno, M., Tonn, R. J., and Reader, P. M. (1972). Studies on the life budget of *Aedes aegypti* in Wat Samphaya, Bangkok, Thailand. Bulletin of the World Health Organization, 46(2):211–226.

[64] Taylor, C., Touré, Y. T., Carnahan, J., Norris, D. E., Dolo, G., Traoré, S. F., Edillo, F. E., and Lanzaro, G. C. (2001). Gene flow among populations of the malaria vector, anopheles gambiae, in mali, west africa. Genetics, 157(2):743–750.

[65] Teixeira, L., Ferreira, A., and Ashburner, M. (2008). The bacterial symbiont Wolbachia induces resistance to RNA viral infections in Drosophila melanogaster. PLOS Biology, 6(12):e2.

[66] Thomson, M. C., Connor, S. J., Quinones, M. L., Jawara, M., Todd, J., and Greenwood, B. M. (1995). Movement of Anopheles gambiae s.l. malaria vectors between villages in The Gambia. Medical and Veterinary Entomology, 9(4):413–419.

[67] Touré, Y. T., Dolo, G., Petrarca, V., Traoré, Bouaré, Dao, A., Carnahan, J., and Taylor, C. E. (1998). Mark–release–recapture experiments with Anopheles gambiae s.l. in Banambani Village, Mali, to determine population size and structure. Medical and Veterinary Entomology, 12(1):74–83.

[68] Unckless, R. L., Clark, A. G., and Messer, P. W. (2017). Evolution of Resistance Against CRISPR/Cas9 Gene Drive. Genetics, 205(2):827–841.

[69] WHO (2017). Global Vector Control Response 2017–2030. World Health Organization, Geneva.

[70] WHO (2017). Technical advisory meeting on risk assessment frameworks for genetically modified *Aedes aegypti* mosquitoes. Venue: HQ, Geneva, Switzerland.

[71] WHO/TDR and FNIH (2014). The guidance framework for testing genetically modified mosquitoes. pp159.

[72] Williams, C. R., Johnson, P. H., Long, S. A., Rapley, L. P., and Ritchie, S. A. (2008). Rapid estimation of Aedes aegypti population size using simulation modeling, with a novel approach to calibration and field validation. Journal of Medical Entomology, 45(6):1173–1179.

[73] World Bank (2016). The short-term economic costs of zika in latin american and the caribbean. Accessed: 28-April-2018.

[74] Yakob, L., Alphey, L., and Bonsall, M. B. (2008). Aedes aegypti control: the concomitant role of competition, space and transgenic technologies. Journal of Applied Ecology, 45(4):1258–1265.

[75] Yakob, L. and Yan, G. (2010). A network population model of the dynamics and control of African malaria vectors. Transactions of The Royal Society of Tropical Medicine and Hygiene, 104(10):669–675.

[76] Zhang, X. H., Tee, L. Y., Wang, X. G., Huang, Q. S., and Yang, S. H. (2015). Off-target Effects in CRISPR/Cas9-mediated Genome Engineering. Molecular Therapy Nucleic Acids, 4:e264.

