## Appendices for "The effect of dispersal and preferential mating on the genetic control of mosquitoes"

### Supporting information

#### S1 Details of ecological model

Parameter values for the population model are listed in table 1. The spatial structure of the network within which all simulations are run, an example of which is shown in fig. 14, is created anew for each simulation by generating fifty coordinate pairs for each village using a normal distribution centred in the middle of three quadrants of the domain. The genetic controls are only released in Village 1 (bottom left quadrant). Village 2 (top left quadrant) represents a nearby settlement; Village 3 (top right quadrant) represents a settlement that is relatively far from Village 1.

##### S1a Mosquito movement

Migration is governed by a weighting matrix $\sigma$, whose elements are Laplace-type dispersal kernels

$$\sigma_{,kl} = \frac{\zeta(d_{,kl})}{Z_{,l}} \tag{S1.1}$$

where

$$\zeta(d_{,kl}) = \omega_1 + \frac{1}{2}\mathrm{e}^{-|d_{,kl}|/\omega_2}, \tag{S1.2}$$

and $Z_{,l} = \sum_{r\neq l}\zeta(d_{rl})$, meaning each column of $\sigma$ sums to unity but each row does not. The distance between nodes $k$ and $l$ is $d_{,kl}$ (thus $d_{kl} = d_{lk}$). A matrix $\bar{P}_{,kl}$, which governs which node connections are open, is constructed by comparing the elements of a lower triangular matrix $P$, where

$$P_{,kl} = \omega_3 + \frac{K}{\sqrt{2\pi D^2}}\mathrm{e}^{-\frac{d^2_{,kl}}{2\omega_4^2 D^2}}, \qquad k > l, \tag{S1.3}$$

to a lower triangular matrix of random numbers between zero and one drawn from a uniform distribution, $U(0,1)$. For $k > l$, if $P_{,kl} \geq U(0,1)_{,kl}$, then $\bar{P}_{,kl} = \bar{P}_{,lk} = 1$; else, $\bar{P}_{,kl} = \bar{P}_{,lk} = 0$. Diagonal elements of $\bar{P}$ are zero. In (S1.3), $K$ defines the average number of connections per node and $D$ defines the average length of the connections. The scale parameters in (S1.3) are set at $\omega_3 = 0.01$ $\omega_4 = 8$ to reproduce reported movement behaviours [32, 34, 64, 66]. The matrix $U(0,1)$ is freshly generated each day such that connections can open and close between nodes through time to simulate physical phenomenon that prevent movement, such as heavy rainfall or closed windows. Element-wise multiplication of $\sigma$ and $\bar{P}$ creates a new matrix which is identical to $\sigma$ except some elements have been set to zero due to the node connections being switched off. The matrix $\bar{\sigma}$ in (2.1) is formed from the element-wise combination of $\sigma$ and $\bar{P}$ by renormalizing each column to sum to one.

Example dispersal behaviours over a single lifetime (roughly thirty days) of the two dispersal archetypes, *Anopheles* and *Aedes*, are shown in fig. 15. Both species have the same $P_{,kl}$ distribution, but their Laplace kernels $\zeta(d_{k,l})$ are different: for *Aedes*, the scale parameters in (S1.2) take the values $\omega_1 = 5 \times 10^{-5}$,

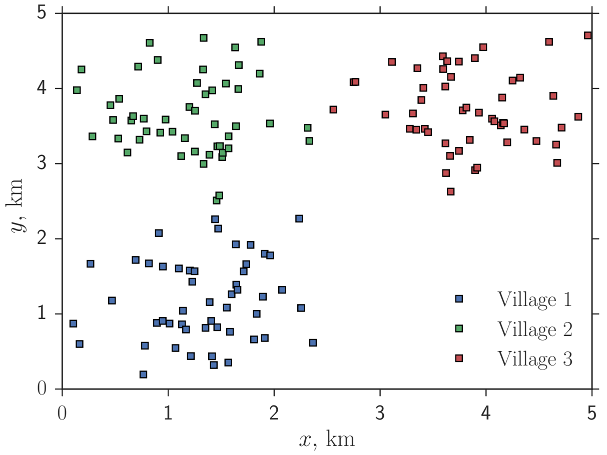


**Figure 14** An example of the spatial distribution of nodes in a 5km domain, constituting three colour-coded villages. Releases of genetically modified mosquitoes are made in Village 1 (bottom left quadrant). Each node represents an area containing a small number of larval habitats, for example a house within the village. Connections between nodes open and close stochastically each day, with probability varying with distance as governed by (S1.3).

$\omega_2 = 0.75$; for *Anopheles*, $\omega_1 = 0.02$, $\omega_2 = 12.5$. These values were chosen to roughly simulate daily dispersal behaviours wherein approximately 99% of migrants stay within 395m and 1.25km of their starting positions, for *Aedes* and *Anopheles* respectively [32, 34, 64, 66]. Additionally, the proportion of a node's population migrating each day is randomly sampled from a Gaussian distribution with mean 0.075 (7.5%) and standard deviation 0.0375. Negative values were set to zero.

The dispersal behaviour is tweaked during the dispersal study in section 3.1 by varying the $\omega_1$ and $\omega_2$ parameters in (S1.2) via the relation

$$\omega_j = \left(\frac{D_l}{0.75}\right)^{n_{\text{scale}}} \omega_j^{(\text{An})}, \tag{S1.4}$$

where

$$n_{\text{scale}} = \frac{\ln\left(\omega_j^{(\text{An})}/\omega_j^{(\text{Ae})}\right)}{\ln 3}, \tag{S1.5}$$

for the *Aedes* and *Anopheles* values of the parameter, $\omega_j^{(\text{Ae})}$ and $\omega_j^{(\text{An})}$, respectively. The dispersal level $D_l$ varies between 0 and 1 and takes the value of the parameters $\omega_1, \omega_2$ smoothly between the *Aedes* value at $D_l = 0.25$ and the *Anopheles* value at $D_l = 0.75$. Too, the dispersal level $D_l$ scales the proportion of a node's population migrating each day, shifting the mean of the Gaussian distribution from which the migration percentage is drawn from 0% at $D_l = 0$ to 15% at $D_l = 1$.

#### S1b Seasonality

Each node in the spatial network has an equilibrium vector population (neglecting migration) that depends upon a nominal nodal human population $H$ and a vector-host ratio $k^*$. Then, $N^{(1)*} = k^* H$, and the population regulation enters the dynamic system (2.1) through the scale of the larval density

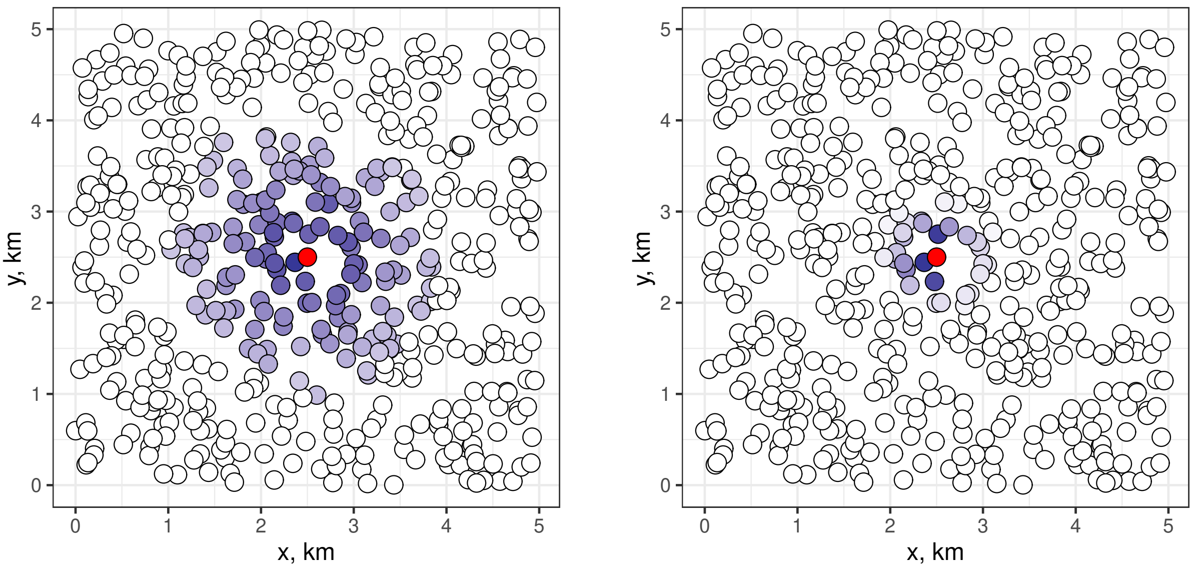


**(a)** *Anopheles* **(b)** *Aedes*

**Figure 15** Dispersal from a single, central populated node (red) into empty nodes after 30 days. Darker nodes are more densely populated. White nodes are empty. (a) Shows *Anopheles* mosquitoes travelling up to around 1.5km in a lifetime, while (b) shows *Aedes* mosquitoes remaining more contained, staying within $\sim$ 500m.

dependence $\nu$ (at a given node). Defining

$$\nu = \frac{m}{2k^* H\mu}\left(\mathrm{e}^{\{\rho m/(2\mu)-m-\mu^B\}}-1\right)^{\frac{1}{\eta}}, \tag{S1.6}$$

we ensure that the wild vector equilibrium population takes the desired value, and we implicitly assume that all genotypes compete equally for resources at the larval stage.

To capture seasonal variation and environmental stochasticity in the carrying capacity of a node, we link the value of $k^*$ to a noisy sinusoid,

$$k^* = 15 + S_a \sin\left(\frac{2\pi t}{365}\right) + \varepsilon_e, \tag{S1.7}$$

where $S_a = 30$ is the amplitude of the seasonal dynamics, to emulate a population that drops to a low level in the dry season, and $\varepsilon_e$ is random normal noise representing localised environmental stochasticity, with a mean of zero and a standard deviation of 12. To prevent very small or negative values of $k^*$, values smaller than 0.1 are tempered by a second definition (the dry/cold season definition):

$$k^* = 0.2 + \varepsilon_1, \tag{S1.8}$$

where $\varepsilon_1$ is random normal noise with a mean of zero and a standard deviation of one. The value of $k^*$ is recalculated for every node each day; an example of its form over three years is shown in fig. 16.

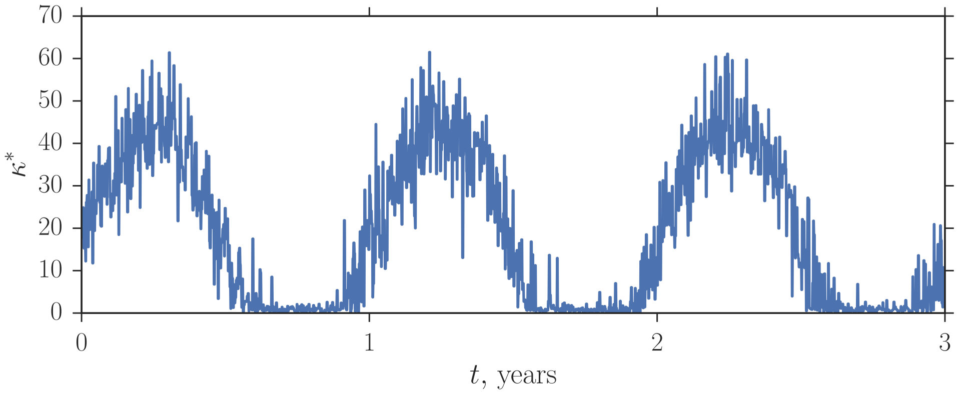


**Figure 16** An example of seasonal dynamics with localised environmental stochasticity of the vector-host ratio $k^*$, which regulates the population dynamics through the definition of the scale of larval density dependence, (S1.6)

### S2 Release strategy

Here we perform several experiments to determine a suitable release strategy with which to carry out the invesigations in the Results section of the main text. We perform these experiments on a single technology, the combined gene drive (CGD) with a homing rate of 0.9 (90% homing efficiency), and with the 'average' mosquito species with a dispersal behaviour of 0.5, half way between the *Aedes* and *Anopheles* archetypes. Initially we use a release strategy wherein mosquitoes are released uniformly at every node in Village 1 at a constant rate of eight per day (400 per day over the whole village) for one year. Results are colour and shape-coded as in the main text, with green (triangles), blue (diamonds) and red (squares) to indicate suppression in Village 1, Villages 1 and 2, and Villages 1, 2 and 3, respectively. Further colour coding indicates whether suppression is temporary (light) or lasting (dark). Gold colouring indicates that suppression was delayed until the year after the control period (for example a gold square denotes suppression in all three villages that takes effect the year after releases cease, while a gold triangle denotes suppression in Village 1 the year after releases cease). Black circles denote simulations wherein no suppression is achieved in any village. Figure 17a shows the results as the lethality efficiency, $c_2$, and the ambient fitness cost, $c_1$, are varied between zero and one. We note two important observations: increasing the lethality efficiency tends to decrease the ability of the control to spread spatially; and sustained suppression can be achieved with relatively low lethality efficiencies if there is a moderate ambient fitness cost (the drive is effectively acting as a mild bi-sex lethal gene in this region of parameter space). If we look at the population timeseries for each village for four subsets of results in fig. 17a defined by the horizontal slices $0 < c_2 < 0.04$, $0.04 < c_2 < 0.08$, $0.08 < c_2 < 0.12$ and $0.12 < c_2 < 0.16$, we see that high lethality efficiency increases the speed with which suppression occurs in Village 1, but decreases the chance of suppression persisting after releases stop or spreading to neighbouring villages. This may be due to repopulation from unsuppressed populations. Considering the above, we choose to set the ambient fitness cost at $c_1 = 0.05$ to model a small negative effect of carrying modified genes, and the lethality efficiency at $c_2 = 0.85$ to allow for technological inefficiencies.

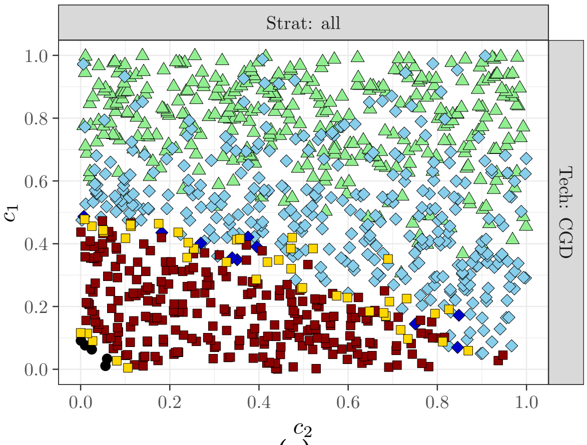

**(a)**

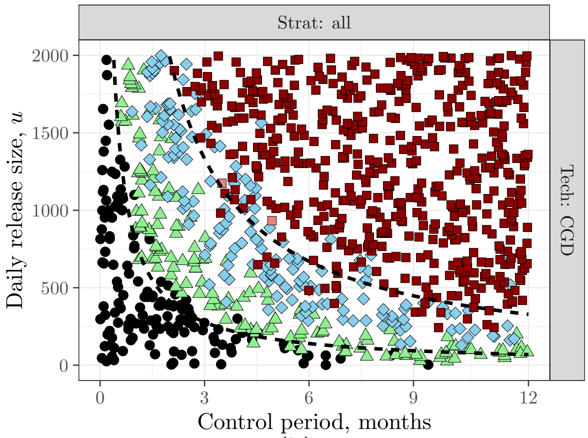

**(b)**

**Figure 17** (a) The effect of lethality efficiency ($c_2$) and ambient fitness cost ($c_1$) on control efforts in thee neighbouring villages. The control strategy is the release of 400 modified males per day for one year, spread over the 50 nodes of Village 1. (b) The effect of release size ($u$) and the length of the control period is examined, for an ambient fitness cost of $c_1 = 0.05$ and a lethality efficiency of $c_2 = 0.85$. In both plots, a combined gene drive (CGD) technology is used. Green (triangles), blue (diamonds) and red (squares) indicate suppression in Village 1, Villages 1 and 2, and Villages 1, 2 and 3, respectively. Further colour coding indicates whether suppression is temporary (light) or lasting (dark). Gold colouring indicates that suppression was delayed until the year after the control period (for example a gold square denotes suppression in all three villages that takes effect the year after releases cease, while a gold triangle denotes suppression in Village 1 the year after releases cease). Black circles denote simulations wherein no suppression is achieved in any village.

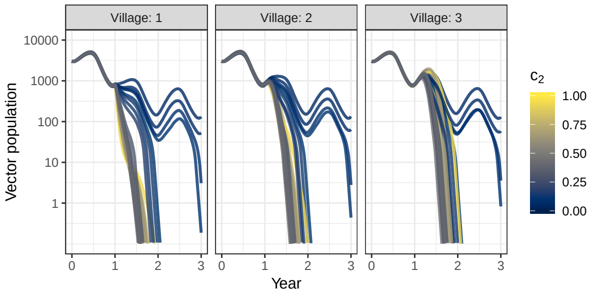

**(a)** $0 < c_1 < 0.04$

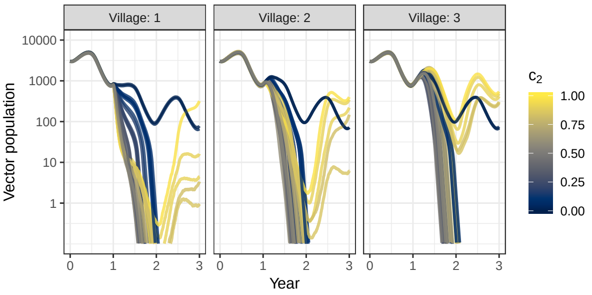

**(b)** $0.04 < c_1 < 0.08$

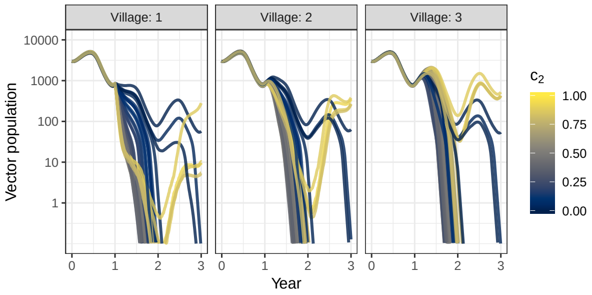

**(c)** $0.08 < c_1 < 0.12$

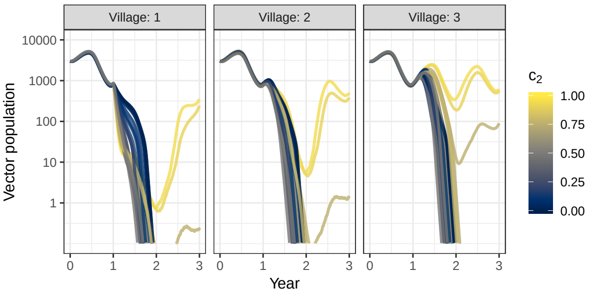

**(d)** $0.12 < c_1 < 0.16$

**Figure 18** Time series showing how the total vector population in each village varies before, during and after the control period, for the simulation results plotted in fig. 17a. The four subplots show time series for the subset of the ambient fitness cost indicated, which translates to thin horizontal strips across fig. 17a. Results are colour coded by the value of the lethality efficiency ($c_2$) for that simulation.

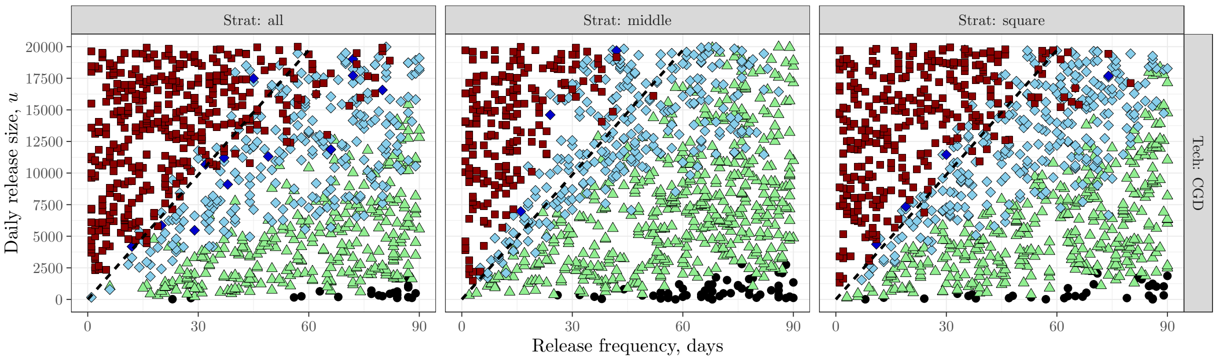


**Figure 19** Control results for pulsed releases of varying sizes and frequencies, with three different spatial release distributions in Village 1: "all", which spreads the release over all nodes in the village; "middle", which concentrates the release at a single node at the centre of the village; and "square", which divides the release up into four equidistant nodes that form a square around the centre of the village, equally spaced betweem the centre and the village edge. The ambient fitness cost is $c_1 = 0.05$ and the lethality efficiency is $c_2 = 0.85$, and a combined gene drive (CGD) technology is used. Colour and marker shape coding as in fig. 17.

We investigate whether a strategy of larger releases over a shorter time or smaller releases over a longer time lead to the most efficient and effective suppression in fig. 17b. The two dashed lines, representing contours of constant total release size (25000 and 120000 insects), demonstrate that using the same number of mosquitoes over a longer period produces marginally better suppression. Thus, we choose a year-long control period for the simulations in the main text.

Because releasing at every house or mosquito breeding spot in a village every day for a year is unfeasible, we examine alternate release strategies – releasing at a single node in the middle of the village ("middle") and releasing at four positions in a square, equally spaced between the middle and the edge of the village ("square") – and release schedules – pulsed releases with a variety of sizes and frequencies (fig. 19). The dashed line in each subplot represents a contour of constant total release size (120000 insects); we see that the square strategy is the most successful at reproducing the performance of the blanket releases ("all"). Bigger, infrequent pulses appear to be more susceptible to stochasticity than smaller, more frequent pulses (fig. 19), so we choose to release weekly pulses of 2000 modified male mosquitoes, spread over the four nodes defined in the square release strategy. This size of release is well within the realms of possibility when the capacity production of large insect rearing facilities can be many millions per month.

### S3 Genotype frequencies

Here we show the proportions of offspring carrying each genotype for the three control technologies tested in the main text, and show how we capture the mating success rate of genetically modified mosquitoes. The number of adult females and males of genotype $i$ in a given node are given by $N^{(i)}$ and $\hat{N}^{(i)}$, respectively; the total number of females and males in a given node are $N^f$ and $N^m$, respectively:

$$N^f = \sum_{i=0}^{n_g} N^{(i)}, \qquad\qquad N^m = \sum_{i=0}^{n_g} \hat{N}^{(i)}. \tag{S3.1}$$

First we discuss how wild-type mating choice is modelled.

#### S3a Wild-type mating preference

Consider the simple case of a population containing one wild-type allele, $A$, and one mutant allele $a$. There are three possible genotypes $AA$, $Aa$ and $aa$, where $AA$ is the wild type. If random mating is assumed between all genotypes, the proportions $p^{(i)}$ of offspring (sired at the current time point) belonging to each genotype are

$$\begin{aligned}
p^{(AA)} &= \frac{1}{2N^m N^f}\left(N^{(AA)}\hat{N}^{(AA)} + \frac{1}{2}\left(N^{(AA)}\hat{N}^{(Aa)} + N^{(Aa)}\hat{N}^{(AA)}\right) + \frac{1}{4}N^{(Aa)}\hat{N}^{(Aa)}\right), && \text{(S3.2a)}\\
p^{(Aa)} &= \frac{1}{2N^m N^f}\left(\frac{1}{2}\left(N^{(AA)}\hat{N}^{(Aa)} + N^{(Aa)}\hat{N}^{(AA)}\right) + N^{(AA)}\hat{N}^{(aa)} + N^{(aa)}\hat{N}^{(AA)}\right.\\
&\qquad \left. + \frac{1}{2}N^{(Aa)}\hat{N}^{(Aa)} + \frac{1}{2}\left(N^{(Aa)}\hat{N}^{(aa)} + N^{(aa)}\hat{N}^{(Aa)}\right)\right), && \text{(S3.2b)}\\
p^{(aa)} &= \frac{1}{2N^m N^f}\left(\frac{1}{4}N^{(Aa)}\hat{N}^{(Aa)} + \frac{1}{2}\left(N^{(Aa)}\hat{N}^{(aa)} + N^{(aa)}\hat{N}^{(Aa)}\right) + N^{(aa)}\hat{N}^{(aa)}\right). && \text{(S3.2c)}
\end{aligned}$$

Now, suppose a fraction $0 \leq \xi \leq 1$ of the wild-type females (always $N^{(1)}$) from each encounter with non-wild males choose instead to mate with wild-type males (always $\hat{N}^{(1)}$). The number of matings between wild-type males and females increases by a related proportion $\chi$. Altering (S3.2) to account for these changes, we find

$$\begin{aligned}
\bar{p}^{(AA)} &= \frac{1}{N^m N^f}\left((1+\chi)N^{(AA)}\hat{N}^{(AA)} + \frac{1}{2}\left((1-\xi)N^{(AA)}\hat{N}^{(Aa)} + N^{(Aa)}\hat{N}^{(AA)}\right)\right.\\
&\qquad \left. + \frac{1}{4}N^{(Aa)}\hat{N}^{(Aa)}\right), && \text{(S3.3a)}\\
\bar{p}^{(Aa)} &= \frac{1}{N^m N^f}\left(\frac{1}{2}\left((1-\xi)N^{(AA)}\hat{N}^{(Aa)} + N^{(Aa)}\hat{N}^{(AA)}\right) + (1-\xi)N^{(AA)}\hat{N}^{(aa)}\right.\\
&\qquad \left. + N^{(aa)}\hat{N}^{(AA)} + \frac{1}{2}N^{(Aa)}\hat{N}^{(Aa)} + \frac{1}{2}\left(N^{(Aa)}\hat{N}^{(aa)} + N^{(aa)}\hat{N}^{(Aa)}\right)\right), && \text{(S3.3b)}\\
\bar{p}^{(aa)} &= \frac{1}{N^m N^f}\left(\frac{1}{4}N^{(Aa)}\hat{N}^{(Aa)} + \frac{1}{2}\left(N^{(Aa)}\hat{N}^{(aa)} + N^{(aa)}\hat{N}^{(Aa)}\right) + N^{(aa)}\hat{N}^{(aa)}\right). && \text{(S3.3c)}
\end{aligned}$$

Due to the mating preference displayed by the wild-type females, we logically expect $\bar{p}^{(AA)} > p^{(AA)}$, $\bar{p}^{(Aa)} < p^{(Aa)}$ and $\bar{p}^{(aa)} = p^{(aa)}$. The constraint

$$\sum_{i=0}^{n_g} p^{(i)} = 1 = \sum_{i=0}^{n_g} \bar{p}^{(i)} \tag{S3.4}$$

must also still hold. Calculating the sums (S3.4) and then subtracting one from the other we find an expression for $\chi$ in terms of $\xi$:

$$\chi = \frac{\frac{1}{2}\hat{N}^{(Aa)} + \frac{1}{2}\hat{N}^{(Aa)} + \hat{N}^{(aa)}}{\hat{N}^{(AA)}}\,\xi, \tag{S3.5}$$

where the fraction is simply the sum of males in non-preferred matings over the number of males in the preferred mating. Thus we may use (S3.5) to define a general expression for the mating success parameter $\chi$, for a given $\xi$, as

$$\chi = \frac{\sum_{i=2}^{n_g} \hat{N}^{(i)}}{\hat{N}^{(1)}}\,\xi. \tag{S3.6}$$

If the wild females struggle to find wild males to mate with, for instance if the wild male population is only a small fraction of the total male population, the females' ability to enact mate choosiness may decrease. We capture this is our model by replacing $\xi$ with $\bar{\xi}$ when $\hat{N}^{(1)}/N^m < 0.25$:

$$\bar{\xi} = 4\frac{\hat{N}^{(1)}}{N^m}\xi. \tag{S3.7}$$

This scales smoothly to $\bar{\xi} = 0$ when there are no wild males left at a given node, meaning the wild females have no choice but to mate normally with the modified males.

#### S3b Self-limiting lethal gene

For the self-limiting lethal gene technology, the wild-type allele is $A$, and released males are homozygote for an engineered construct $H$. The offspring proportions $p^{(1)}$, $p^{(2)}$ and $p^{(3)}$ relate to genotypes $AA$, $AH$ and $HH$, respectively. These proportions are given by

$$p^{(1)} = \frac{1}{2N^m N^f}\left((1+\chi)N^{(1)}\hat{N}^{(1)} + \frac{1}{2}((1-\xi)N^{(1)}\hat{N}^{(2)} + \hat{N}^{(1)}N^{(2)}) + \frac{1}{4}N^{(2)}\hat{N}^{(2)}\right), \tag{S3.8a}$$

$$p^{(2)} = \frac{1}{2N^m N^f}\left(\frac{1}{2}((1-\xi)N^{(1)}\hat{N}^{(2)} + \hat{N}^{(1)}N^{(2)}) + ((1-\xi)N^{(1)}\hat{N}^{(3)} + \hat{N}^{(1)}N^{(3)}) + \frac{1}{2}N^{(2)}\hat{N}^{(2)} + \frac{1}{2}(N^{(2)}\hat{N}^{(3)} + \hat{N}^{(2)}N^{(3)})\right), \tag{S3.8b}$$

$$p^{(3)} = \frac{1}{2N^m N^f}\left(\frac{1}{4}N^{(2)}\hat{N}^{(2)} + \frac{1}{2}(N^{(2)}\hat{N}^{(3)} + \hat{N}^{(2)}N^{(3)}) + N^{(3)}\hat{N}^{(3)}\right). \tag{S3.8c}$$

The female-specific lethal gene $H$ has a heterozygote lethality efficiency of 0.85, and any mosquito (male or female) carrying the allele $H$ is struck by an ambient fitness cost of 0.05. Fitness costs combine multiplicatively.

#### S3c Homologous underdominance threshold drive

The homologous underdominance model has a wild-type allele $A$ (and wild-type genotype $AA$) and engineered underdominant constructs $\alpha$ and $\beta$ that both express a lethal toxin and suppress the toxin expression by the other engineered allele (thus $A\alpha$ and $A\beta$ suffer imposed mortality while $\alpha\beta$ suffers only ambient fitness costs). The offspring proportions $p^{(1)}$, $p^{(2)}$, $p^{(3)}$, $p^{(4)}$, $p^{(5)}$, $p^{(6)}$ relate to genotypes $AA$,

$A\alpha$, $A\beta$, $\alpha\beta$, $\alpha\alpha$ and $\beta\beta$, respectively. These offspring proportions are

$$\begin{aligned}
p^{(1)} =& \frac{1}{2N_m N_f}\bigg((1+\chi)N^{(1)}\hat{N}^{(1)} + \frac{1}{2}((1-\xi)N^{(1)}\hat{N}^{(2)} + \hat{N}^{(1)}N^{(2)}) + \frac{1}{4}N^{(2)}\hat{N}^{(2)} \\
&+\frac{1}{2}((1-\xi)N^{(1)}\hat{N}^{(3)} + \hat{N}^{(1)}N^{(3)}) + \frac{1}{4}N^{(3)}\hat{N}^{(3)} + \frac{1}{4}(N^{(2)}\hat{N}^{(3)} + \hat{N}^{(2)}N^{(3)})\bigg), \qquad \text{(S3.9a)}
\end{aligned}$$

$$\begin{aligned}
p^{(2)} =& \frac{1}{2N_m N_f}\bigg(\frac{1}{2}((1-\xi)N^{(1)}\hat{N}^{(2)} + \hat{N}^{(1)}N^{(2)}) + \frac{1}{2}(\hat{N}^{(1)}N^{(4)} + (1-\xi)N^{(1)}\hat{N}^{(4)}) \\
&+(\hat{N}^{(1)}N^{(5)} + (1-\xi)N^{(1)}\hat{N}^{(5)}) + \frac{1}{2}N^{(2)}\hat{N}^{(2)} + \frac{1}{4}(N^{(2)}\hat{N}^{(3)} + \hat{N}^{(2)}N^{(3)}) \\
&+\frac{1}{4}(\hat{N}^{(2)}N^{(4)} + N^{(2)}\hat{N}^{(4)}) + \frac{1}{2}(\hat{N}^{(2)}N^{(5)} + N^{(2)}\hat{N}^{(5)}) + \frac{1}{2}(N^{(3)}\hat{N}^{(5)} + \hat{N}^{(3)}N^{(5)}) \\
&+\frac{1}{4}(\hat{N}^{(3)}N^{(4)} + N^{(3)}\hat{N}^{(4)})\bigg), \qquad \text{(S3.9b)}
\end{aligned}$$

$$\begin{aligned}
p^{(3)} =& \frac{1}{2N_m N_f}\bigg(\frac{1}{2}((1-\xi)N^{(1)}\hat{N}^{(3)} + \hat{N}^{(1)}N^{(3)}) + \frac{1}{2}(\hat{N}^{(1)}N^{(4)} + (1-\xi)N^{(1)}\hat{N}^{(4)}) \\
&+((1-\xi)N^{(1)}\hat{N}^{(6)} + \hat{N}^{(1)}N^{(6)}) + \frac{1}{2}(N^{(2)}\hat{N}^{(6)} + \hat{N}^{(2)}N^{(6)}) + \frac{1}{2}N^{(3)}\hat{N}^{(3)} \\
&+\frac{1}{4}(N^{(2)}\hat{N}^{(3)} + \hat{N}^{(2)}N^{(3)}) + \frac{1}{4}(\hat{N}^{(2)}N^{(4)} + N^{(2)}\hat{N}^{(4)}) \\
&+\frac{1}{2}(N^{(3)}\hat{N}^{(6)} + \hat{N}^{(3)}N^{(6)}) + \frac{1}{4}(\hat{N}^{(3)}N^{(4)} + N^{(3)}\hat{N}^{(4)})\bigg), \qquad \text{(S3.9c)}
\end{aligned}$$

$$\begin{aligned}
p^{(4)} =& \frac{1}{2N_m N_f}\bigg(\frac{1}{4}(N^{(2)}\hat{N}^{(3)} + \hat{N}^{(2)}N^{(3)}) + \frac{1}{4}(\hat{N}^{(2)}N^{(4)} + N^{(2)}\hat{N}^{(4)}) \\
&+\frac{1}{2}(N^{(2)}\hat{N}^{(6)} + \hat{N}^{(2)}N^{(6)}) + \frac{1}{4}(\hat{N}^{(3)}N^{(4)} + N^{(3)}\hat{N}^{(4)}) + \frac{1}{2}N^{(4)}\hat{N}^{(4)} \\
&+\frac{1}{2}(N^{(3)}\hat{N}^{(5)} + \hat{N}^{(3)}N^{(5)}) + (N^{(5)}\hat{N}^{(6)} + \hat{N}^{(5)}N^{(6)}) + \frac{1}{2}(\hat{N}^{(5)}N^{(4)} + N^{(5)}\hat{N}^{(4)}) \\
&+\frac{1}{2}(\hat{N}^{(6)}N^{(4)} + N^{(6)}\hat{N}^{(4)})\bigg), \qquad \text{(S3.9d)}
\end{aligned}$$

$$\begin{aligned}
p^{(5)} =& \frac{1}{2N_m N_f}\bigg(\frac{1}{4}N^{(2)}\hat{N}^{(2)} + \frac{1}{4}(\hat{N}^{(2)}N^{(4)} + N^{(2)}\hat{N}^{(4)}) + \frac{1}{2}(N^{(2)}\hat{N}^{(5)} + \hat{N}^{(2)}N^{(5)}) \\
&+\frac{1}{4}N^{(4)}\hat{N}^{(4)} + \frac{1}{2}(\hat{N}^{(5)}N^{(4)} + N^{(5)}\hat{N}^{(4)}) + N^{(5)}\hat{N}^{(5)}\bigg), \qquad \text{(S3.9e)}
\end{aligned}$$

$$\begin{aligned}
p^{(6)} =& \frac{1}{2N_m N_f}\bigg(\frac{1}{4}N^{(3)}\hat{N}^{(3)} + \frac{1}{4}(\hat{N}^{(3)}N^{(4)} + N^{(3)}\hat{N}^{(4)}) + \frac{1}{2}(N^{(3)}\hat{N}^{(6)} + \hat{N}^{(3)}N^{(6)}) \\
&+\frac{1}{4}N^{(4)}\hat{N}^{(4)} + \frac{1}{2}(\hat{N}^{(6)}N^{(4)} + N^{(6)}\hat{N}^{(4)}) + N^{(6)}\hat{N}^{(6)}\bigg). \qquad \text{(S3.9f)}
\end{aligned}$$

Modified males of genotype $\alpha\beta$ are released. The toxic fitness cost imposed by a single construct is female-specific and has a lethality efficiency of 0.85. Any mosquito (male or female) carrying a modified allele $\alpha$, $\beta$ is struck with an ambient relative fitness cost of 0.05. Fitness costs combine multiplicatively.

#### S3d Non-homologous underdominance threshold drive

The non-homologous underdominance model has wild-type alleles $A$ and $B$ (and wild-type genotype $AABB$) and engineered underdominant constructs $\alpha$ and $\beta$ that both express a lethal toxin and suppress the toxin expression by the other engineered allele (thus $\alpha\alpha BB$ and $AA\beta\beta$ suffer imposed mortality while $\alpha\alpha\beta\beta$ suffers only unintended fitness costs). The offspring proportions $p^{(1)}$, $p^{(2)}$, $p^{(3)}$, $p^{(4)}$, $p^{(5)}$,

$p^{(6)}$, $p^{(7)}$, $p^{(8)}$ and $p^{(9)}$ relate to genotypes $AABB$, $A\alpha BB$, $AAB\beta$, $A\alpha B\beta$, $\alpha\alpha BB$, $AA\beta\beta$, $\alpha\alpha B\beta$, $A\alpha\beta\beta$ and $\alpha\alpha\beta\beta$, respectively. The offspring proportions are

$$\begin{aligned}
p^{(1)} = \frac{1}{2N^mN^f} \Big( & (1+\chi)N^{(1)}\hat{N}^{(1)} + \frac{1}{2}((1-\xi)N^{(1)}\hat{N}^{(2)} + \hat{N}^{(1)}N^{(2)}) + \frac{1}{4}N^{(2)}\hat{N}^{(2)} \\
& + \frac{1}{2}((1-\xi)N^{(1)}\hat{N}^{(3)} + \hat{N}^{(1)}N^{(3)}) + \frac{1}{4}((1-\xi)N^{(1)}\hat{N}^{(4)} + \hat{N}^{(1)}N^{(4)}) \\
& + \frac{1}{4}(N^{(2)}\hat{N}^{(3)} + \hat{N}^{(2)}N^{(3)}) + \frac{1}{16}N^{(4)}\hat{N}^{(4)} + \frac{1}{8}(N^{(2)}\hat{N}^{(4)} + \hat{N}^{(2)}N^{(4)}) \\
& + \frac{1}{4}N^{(3)}\hat{N}^{(3)} + \frac{1}{8}(N^{(3)}\hat{N}^{(4)} + \hat{N}^{(3)}N^{(4)})\Big), \qquad \text{(S3.10a)}
\end{aligned}$$

$$\begin{aligned}
p^{(2)} = \frac{1}{2N^mN^f} \Big( & \frac{1}{2}((1-\xi)N^{(1)}\hat{N}^{(2)} + \hat{N}^{(1)}N^{(2)}) + \frac{1}{4}((1-\xi)N^{(1)}\hat{N}^{(4)} + \hat{N}^{(1)}N^{(4)}) \\
& + ((1-\xi)N^{(1)}\hat{N}^{(5)} + \hat{N}^{(1)}N^{(5)}) + \frac{1}{2}((1-\xi)N^{(1)}\hat{N}^{(7)} + \hat{N}^{(1)}N^{(7)}) + \frac{1}{2}N^{(2)}\hat{N}^{(2)} \\
& + \frac{1}{4}(N^{(2)}\hat{N}^{(3)} + \hat{N}^{(2)}N^{(3)}) + \frac{1}{4}(N^{(2)}\hat{N}^{(4)} + \hat{N}^{(2)}N^{(4)}) + \frac{1}{2}(N^{(2)}\hat{N}^{(5)} + \hat{N}^{(2)}N^{(5)}) \\
& + \frac{1}{4}(N^{(2)}\hat{N}^{(7)} + \hat{N}^{(2)}N^{(7)}) + \frac{1}{8}(N^{(3)}\hat{N}^{(4)} + \hat{N}^{(3)}N^{(4)}) + \frac{1}{8}N^{(4)}\hat{N}^{(4)} \\
& + \frac{1}{2}(N^{(3)}\hat{N}^{(5)} + \hat{N}^{(3)}N^{(5)}) + \frac{1}{4}(N^{(3)}\hat{N}^{(7)} + \hat{N}^{(3)}N^{(7)}) + \frac{1}{4}(N^{(4)}\hat{N}^{(5)} + \hat{N}^{(4)}N^{(5)}) \\
& + \frac{1}{8}(N^{(4)}\hat{N}^{(7)} + \hat{N}^{(4)}N^{(7)})\Big), \qquad \text{(S3.10b)}
\end{aligned}$$

$$\begin{aligned}
p^{(3)} = \frac{1}{2N^mN^f} \Big( & \frac{1}{2}((1-\xi)N^{(1)}\hat{N}^{(3)} + \hat{N}^{(1)}N^{(3)}) + \frac{1}{4}((1-\xi)N^{(1)}\hat{N}^{(4)} + \hat{N}^{(1)}N^{(4)}) \\
& + ((1-\xi)N^{(1)}\hat{N}^{(6)} + \hat{N}^{(1)}N^{(6)}) + \frac{1}{2}((1-\xi)N^{(1)}\hat{N}^{(8)} + \hat{N}^{(1)}N^{(8)}) + \frac{1}{2}N^{(3)}\hat{N}^{(3)} \\
& + \frac{1}{4}(N^{(2)}\hat{N}^{(3)} + \hat{N}^{(2)}N^{(3)}) + \frac{1}{8}(N^{(2)}\hat{N}^{(4)} + \hat{N}^{(2)}N^{(4)}) + \frac{1}{2}(N^{(2)}\hat{N}^{(6)} + \hat{N}^{(2)}N^{(6)}) \\
& + \frac{1}{4}(N^{(2)}\hat{N}^{(8)} + \hat{N}^{(2)}N^{(8)}) + \frac{1}{4}(N^{(3)}\hat{N}^{(4)} + \hat{N}^{(3)}N^{(4)}) + \frac{1}{2}(N^{(3)}\hat{N}^{(6)} + \hat{N}^{(3)}N^{(6)}) \\
& + \frac{1}{4}(N^{(3)}\hat{N}^{(8)} + \hat{N}^{(3)}N^{(8)}) + \frac{1}{8}N^{(4)}\hat{N}^{(4)} + \frac{1}{4}(N^{(4)}\hat{N}^{(6)} + \hat{N}^{(4)}N^{(6)}) \\
& + \frac{1}{8}(N^{(4)}\hat{N}^{(8)} + \hat{N}^{(4)}N^{(8)})\Big), \qquad \text{(S3.10c)}
\end{aligned}$$

$$\begin{aligned}
p^{(4)} = \frac{1}{2N^mN^f} \Big( & \frac{1}{4}((1-\xi)N^{(1)}\hat{N}^{(4)} + \hat{N}^{(1)}N^{(4)}) + \frac{1}{2}((1-\xi)N^{(1)}\hat{N}^{(7)} + \hat{N}^{(1)}N^{(7)}) \\
& + \frac{1}{2}((1-\xi)N^{(1)}\hat{N}^{(8)} + \hat{N}^{(1)}N^{(8)}) + ((1-\xi)N^{(1)}\hat{N}^{(9)} + \hat{N}^{(1)}N^{(9)}) + \frac{1}{4}N^{(4)}\hat{N}^{(4)} \\
& + \frac{1}{4}(N^{(2)}\hat{N}^{(3)} + \hat{N}^{(2)}N^{(3)}) + \frac{1}{4}(N^{(2)}\hat{N}^{(4)} + \hat{N}^{(2)}N^{(4)}) + \frac{1}{2}(N^{(2)}\hat{N}^{(6)} + \hat{N}^{(2)}N^{(6)}) \\
& + \frac{1}{4}(N^{(2)}\hat{N}^{(7)} + \hat{N}^{(2)}N^{(7)}) + \frac{1}{2}(N^{(2)}\hat{N}^{(8)} + \hat{N}^{(2)}N^{(8)}) + \frac{1}{2}(N^{(2)}\hat{N}^{(9)} + \hat{N}^{(2)}N^{(9)}) \\
& + \frac{1}{4}(N^{(3)}\hat{N}^{(4)} + \hat{N}^{(3)}N^{(4)}) + \frac{1}{2}(N^{(3)}\hat{N}^{(5)} + \hat{N}^{(3)}N^{(5)}) + \frac{1}{2}(N^{(3)}\hat{N}^{(7)} + \hat{N}^{(3)}N^{(7)}) \\
& + \frac{1}{4}(N^{(3)}\hat{N}^{(8)} + \hat{N}^{(3)}N^{(8)}) + \frac{1}{2}(N^{(3)}\hat{N}^{(9)} + \hat{N}^{(3)}N^{(9)}) + \frac{1}{4}(N^{(4)}\hat{N}^{(5)} + \hat{N}^{(4)}N^{(5)}) \\
& + \frac{1}{4}(N^{(4)}\hat{N}^{(6)} + \hat{N}^{(4)}N^{(6)}) + \frac{1}{4}(N^{(4)}\hat{N}^{(7)} + \hat{N}^{(4)}N^{(7)}) + \frac{1}{4}(N^{(4)}\hat{N}^{(8)} + \hat{N}^{(4)}N^{(8)}) \\
& + \frac{1}{4}(N^{(4)}\hat{N}^{(9)} + \hat{N}^{(4)}N^{(9)}) + (N^{(5)}\hat{N}^{(6)} + \hat{N}^{(5)}N^{(6)}) + \frac{1}{2}(N^{(5)}\hat{N}^{(8)} + \hat{N}^{(5)}N^{(8)}) \\
& + \frac{1}{2}(N^{(6)}\hat{N}^{(7)} + \hat{N}^{(6)}N^{(7)}) + \frac{1}{4}(N^{(7)}\hat{N}^{(8)} + \hat{N}^{(7)}N^{(8)})\Big), \qquad \text{(S3.10d)}
\end{aligned}$$

$$p^{(5)} = \frac{1}{2N^mN^f} \Big( \frac{1}{4}N^{(2)}\hat{N}^{(2)} + \frac{1}{8}(N^{(2)}\hat{N}^{(4)} + \hat{N}^{(2)}N^{(4)}) + \frac{1}{2}(N^{(2)}\hat{N}^{(5)} + \hat{N}^{(2)}N^{(5)})$$

$$
\begin{aligned}
&+ \frac{1}{4}(N^{(2)}\hat{N}^{(7)} + \hat{N}^{(2)}N^{(7)}) + \frac{1}{16}N^{(4)}\hat{N}^{(4)} + \frac{1}{4}(N^{(4)}\hat{N}^{(5)} + \hat{N}^{(4)}N^{(5)}) \\
&+ \frac{1}{8}(N^{(4)}\hat{N}^{(7)} + \hat{N}^{(4)}N^{(7)}) + N^{(5)}\hat{N}^{(5)} + \frac{1}{2}(N^{(5)}\hat{N}^{(7)} + \hat{N}^{(5)}N^{(7)}) + \frac{1}{4}N^{(7)}\hat{N}^{(7)}\Bigg), \qquad \text{(S3.10e)}
\end{aligned}
$$

$$
\begin{aligned}
p^{(6)} = \frac{1}{2N^mN^f}\Bigg(&\frac{1}{4}N^{(3)}\hat{N}^{(3)} + \frac{1}{8}(N^{(3)}\hat{N}^{(4)} + \hat{N}^{(3)}N^{(4)}) + \frac{1}{2}(N^{(3)}\hat{N}^{(6)} + \hat{N}^{(3)}N^{(6)}) \\
&+ \frac{1}{4}(N^{(3)}\hat{N}^{(8)} + \hat{N}^{(3)}N^{(8)}) + \frac{1}{16}N^{(4)}\hat{N}^{(4)} + \frac{1}{4}(N^{(4)}\hat{N}^{(6)} + \hat{N}^{(4)}N^{(6)}) + N^{(6)}\hat{N}^{(6)} \\
&+ \frac{1}{8}(N^{(4)}\hat{N}^{(8)} + \hat{N}^{(4)}N^{(8)}) + \frac{1}{2}(N^{(6)}\hat{N}^{(8)} + \hat{N}^{(6)}N^{(8)}) + \frac{1}{4}N^{(8)}\hat{N}^{(8)}\Bigg), \qquad \text{(S3.10f)}
\end{aligned}
$$

$$
\begin{aligned}
p^{(7)} = \frac{1}{2N^mN^f}\Bigg(&\frac{1}{8}(N^{(2)}\hat{N}^{(4)} + \hat{N}^{(2)}N^{(4)}) + \frac{1}{4}(N^{(2)}\hat{N}^{(7)} + \hat{N}^{(2)}N^{(7)}) + \frac{1}{8}N^{(4)}\hat{N}^{(4)} \\
&+ \frac{1}{4}(N^{(2)}\hat{N}^{(8)} + \hat{N}^{(2)}N^{(8)}) + \frac{1}{2}(N^{(2)}\hat{N}^{(9)} + \hat{N}^{(2)}N^{(9)}) + \frac{1}{4}(N^{(4)}\hat{N}^{(5)} + \hat{N}^{(4)}N^{(5)}) \\
&+ \frac{1}{4}(N^{(4)}\hat{N}^{(7)} + \hat{N}^{(4)}N^{(7)}) + \frac{1}{8}(N^{(4)}\hat{N}^{(8)} + \hat{N}^{(4)}N^{(8)}) + \frac{1}{4}(N^{(4)}\hat{N}^{(9)} + \hat{N}^{(4)}N^{(9)}) \\
&+ \frac{1}{2}(N^{(5)}\hat{N}^{(7)} + \hat{N}^{(5)}N^{(7)}) + \frac{1}{2}(N^{(5)}\hat{N}^{(8)} + \hat{N}^{(5)}N^{(8)}) + (N^{(5)}\hat{N}^{(9)} + \hat{N}^{(5)}N^{(9)}) \\
&+ \frac{1}{2}N^{(7)}\hat{N}^{(7)} + \frac{1}{4}(N^{(7)}\hat{N}^{(8)} + \hat{N}^{(7)}N^{(8)}) + \frac{1}{2}(N^{(7)}\hat{N}^{(9)} + \hat{N}^{(7)}N^{(9)})\Bigg), \qquad \text{(S3.10g)}
\end{aligned}
$$

$$
\begin{aligned}
p^{(8)} = \frac{1}{2N^mN^f}\Bigg(&\frac{1}{8}(N^{(3)}\hat{N}^{(4)} + \hat{N}^{(3)}N^{(4)}) + \frac{1}{4}(N^{(3)}\hat{N}^{(7)} + \hat{N}^{(3)}N^{(7)}) + \frac{1}{8}N^{(4)}\hat{N}^{(4)} \\
&+ \frac{1}{4}(N^{(3)}\hat{N}^{(8)} + \hat{N}^{(3)}N^{(8)}) + \frac{1}{2}(N^{(3)}\hat{N}^{(9)} + \hat{N}^{(3)}N^{(9)}) + \frac{1}{4}(N^{(4)}\hat{N}^{(6)} + \hat{N}^{(4)}N^{(6)}) \\
&+ \frac{1}{8}(N^{(4)}\hat{N}^{(7)} + \hat{N}^{(4)}N^{(7)}) + \frac{1}{4}(N^{(4)}\hat{N}^{(8)} + \hat{N}^{(4)}N^{(8)}) + \frac{1}{4}(N^{(4)}\hat{N}^{(9)} + \hat{N}^{(4)}N^{(9)}) \\
&+ \frac{1}{2}(N^{(6)}\hat{N}^{(7)} + \hat{N}^{(6)}N^{(7)}) + \frac{1}{2}(N^{(6)}\hat{N}^{(8)} + \hat{N}^{(6)}N^{(8)}) + (N^{(6)}\hat{N}^{(9)} + \hat{N}^{(6)}N^{(9)}) \\
&+ \frac{1}{2}N^{(8)}\hat{N}^{(8)} + \frac{1}{4}(N^{(7)}\hat{N}^{(8)} + \hat{N}^{(7)}N^{(8)}) + \frac{1}{2}(N^{(8)}\hat{N}^{(9)} + \hat{N}^{(8)}N^{(9)})\Bigg), \qquad \text{(S3.10h)}
\end{aligned}
$$

$$
\begin{aligned}
p^{(9)} = \frac{1}{2N^mN^f}\Bigg(&\frac{1}{16}N^{(4)}\hat{N}^{(4)} + \frac{1}{8}(N^{(4)}\hat{N}^{(7)} + \hat{N}^{(4)}N^{(7)}) + \frac{1}{8}(N^{(4)}\hat{N}^{(8)} + \hat{N}^{(4)}N^{(8)}) \\
&+ \frac{1}{4}(N^{(4)}\hat{N}^{(9)} + \hat{N}^{(4)}N^{(9)}) + \frac{1}{4}N^{(7)}\hat{N}^{(7)} + \frac{1}{4}(N^{(7)}\hat{N}^{(8)} + \hat{N}^{(7)}N^{(8)}) + N^{(9)}\hat{N}^{(9)} \\
&+ \frac{1}{2}(N^{(7)}\hat{N}^{(9)} + \hat{N}^{(7)}N^{(9)}) + \frac{1}{4}N^{(8)}\hat{N}^{(8)} + \frac{1}{2}(N^{(8)}\hat{N}^{(9)} + \hat{N}^{(8)}N^{(9)})\Bigg). \qquad \text{(S3.10i)}
\end{aligned}
$$

Modified males of genotype $\alpha\alpha\beta\beta$ are released. We use a weakly suppressed underdominance system, in which a single $\alpha$ allele cannot suppress the toxin expression of two $\beta$ alleles, such that $A\alpha\beta\beta$ suffers the toxic fitness cost imposed by the $\beta$ allele (and vice versa). The toxic fitness cost is female-specific and has a lethality efficiency of 0.85. Any mosquito (male or female) carrying a modified allele $\alpha$, $\beta$ is struck with an ambient relative fitness cost of 0.05. Fitness costs combine multiplicatively.

#### S3e Combined homing gene drive

The homing gene drive wild-type allele is $A$, and released males are homozygote for the construct $H$. The construct $H$ converts $A$ alleles to $H$ alleles in offspring with homing rate $g$. In all computations the heterozygote homing rate is set at $g = 0.9$, which scales multiplicatively. The offspring proportions $p^{(1)}$,

$p^{(2)}$ and $p^{(3)}$ relate to genotypes $AA$, $AH$ and $HH$, respectively. These proportions are given by

$$p^{(1)} = \frac{1}{2N^m N^f}\Bigg((1+\chi)N^{(1)}\hat{N}^{(1)} + \frac{1}{2}(1-g)((1-\xi)N^{(1)}\hat{N}^{(2)} + \hat{N}^{(1)}N^{(2)}) + \frac{1}{4}(1-g)^2 N^{(2)}\hat{N}^{(2)}\Bigg), \tag{S3.11a}$$

$$p^{(2)} = \frac{1}{2N^m N^f}\Bigg(\frac{1}{2}(1+g)((1-\xi)N^{(1)}\hat{N}^{(2)} + \hat{N}^{(1)}N^{(2)}) + ((1-\xi)N^{(1)}\hat{N}^{(3)} + \hat{N}^{(1)}N^{(3)}) + \frac{1}{2}(1+g)(1-g)N^{(2)}\hat{N}^{(2)} + \frac{1}{2}(1-g)(N^{(2)}\hat{N}^{(3)} + \hat{N}^{(2)}N^{(3)})\Bigg), \tag{S3.11b}$$

$$p^{(3)} = \frac{1}{2N^m N^f}\Bigg(\frac{1}{4}(1+g)^2 N^{(2)}\hat{N}^{(2)} + \frac{1}{2}(1+g)(N^{(2)}\hat{N}^{(3)} + \hat{N}^{(2)}N^{(3)}) + N^{(3)}\hat{N}^{(3)}\Bigg). \tag{S3.11c}$$

The allele $H$ imposes a dominant female-specific lethality, with efficiency 0.85, and any mosquito (male or female) carrying the allele $H$ is struck with an ambient relative fitness cost of 0.05. Fitness costs combine multiplicatively.

#### S3f Separated homing gene drive

In this system the drive allele $D$ (with wild-type allele $w$) is situated at a different locus to the lethal payload allele $V$ (with wild-type allele $U$). The wild-type genotype is $wwUU$ and the released genotype is $DDUU$. The drive allele $D$ causes the allele $V$ to be inserted in place of $U$ with rate of success dependent upon the number of copies of the allele $D$. The heterozygote homing rate is $g_{Dw}$, while the homozygote rate combines as $g_{DD} = 1 - (1 - g_{Dw})^2$. The offspring proportions $p^{(1)}$, $p^{(2)}$, $p^{(3)}$, $p^{(4)}$, $p^{(5)}$, $p^{(6)}$, $p^{(7)}$, $p^{(8)}$ and $p^{(9)}$ relate to genotypes $wwUU$, $DwUU$, $DDUU$, $wwVU$, $DwVU$, $DDVU$, $wwVV$, $DwVV$ and $DDVV$, respectively. The form of the proportions are not given here due to their unsightliness. The default value for the heterozygote homing rate is $g_{Dw} = 0.9$. The allele $V$ imposes a dominant female-specific lethality, with efficiency 0.85, and any mosquito (male or female) carrying a modified allele $D$, $V$ is struck with an ambient relative fitness cost of 0.05. Fitness costs combine multiplicatively.

#### S3g *Wolbachia*

We capture the *Wolbachia* systems in the same population genetic framework as the other technologies, but note that our proportions $p^{(i)}$ no longer refer to genotypes but to infection status. Thus, in the unidirectional *Wolbachia* system, $p^{(1)}$ refers to the proportion of offspring that are uninfected (wild type) and $p^{(2)}$ refers to the infected proportion. Because the action of *Wolbachia* is different to the technologies above, we handle the lethality and drive mechanisms slightly differently. We introduce two new parameters: the leakiness of the infection rate, $\ell \in [0, 1]$, which leads to a small fraction of wild-type offspring emerging from infection-inducing pairings (and can be thought of as one minus the homing rate of a normal gene drive); and the efficiency of the cytoplasmic incompatibility, $\mathrm{CI}_{\mathrm{eff}} \in [0, 1]$, where a value $\mathrm{CI}_{\mathrm{eff}} < 1$ allows offspring to survive that would otherwise have not been viable (and can be thought of as equal to the lethality efficiency of the lethal payload genes in other technologies). The default parameter values are $\ell = 0.1$ and $\mathrm{CI_{eff}} = 0.85$. The offspring proportions for the unidirectional system are

$$p^{(1)} = \frac{1}{2N^m N^f}\left((1+\chi)N^{(1)}\hat{N}^{(1)} + \ell\hat{N}^{(1)}N^{(2)} + \frac{1}{2}(1-\xi)(1-\mathrm{CI_{eff}})N^{(1)}\hat{N}^{(2)}\right), \tag{S3.12a}$$

$$p^{(2)} = \frac{1}{2N^m N^f}\left((1-\ell)\hat{N}^{(1)}N^{(2)} + N^{(2)}\hat{N}^{(2)} + \frac{1}{2}(1-\xi)(1-\mathrm{CI_{eff}})N^{(1)}\hat{N}^{(2)}\right), \tag{S3.12b}$$

while for the bidirectional system they are

$$\begin{aligned} p^{(1)} = \frac{1}{2N^m N^f}\Big((1+\chi)N^{(1)}\hat{N}^{(1)} + \ell\hat{N}^{(1)}N^{(2)} + \ell\hat{N}^{(1)}N^{(3)} \\ + \frac{1}{2}(1-\xi)(1-\mathrm{CI_{eff}})N^{(1)}\hat{N}^{(2)} + \frac{1}{2}(1-\xi)(1-\mathrm{CI_{eff}})N^{(1)}\hat{N}^{(3)}\Big), \end{aligned} \tag{S3.13a}$$

$$\begin{aligned} p^{(2)} = \frac{1}{2N^m N^f}\Big((1-\ell)\hat{N}^{(1)}N^{(2)} + N^{(2)}\hat{N}^{(2)} + \frac{1}{2}(1-\xi)(1-\mathrm{CI_{eff}})N^{(1)}\hat{N}^{(2)} \\ + \frac{1}{2}(1-\mathrm{CI_{eff}})(N^{(2)}\hat{N}^{(3)} + N^{(3)}\hat{N}^{(2)})\Big), \end{aligned} \tag{S3.13b}$$

$$\begin{aligned} p^{(3)} = \frac{1}{2N^m N^f}\Big((1-\ell)\hat{N}^{(1)}N^{(3)} + N^{(3)}\hat{N}^{(3)} + \frac{1}{2}(1-\xi)(1-\mathrm{CI_{eff}})N^{(1)}\hat{N}^{(3)} \\ + \frac{1}{2}(1-\mathrm{CI_{eff}})(N^{(2)}\hat{N}^{(3)} + N^{(3)}\hat{N}^{(2)})\Big), \end{aligned} \tag{S3.13c}$$

where $p^{(3)}$ denotes the proportion of offspring infected with a second strain of the *Wolbachia* bacteria. Any infected mosquito is struck with an ambient fitness cost of 0.05, although we examine the effect of infection causing fitness advantage in section 3.4.
